# NAPE-PLD in the ventral tegmental area regulates reward events, feeding and energy homeostasis

**DOI:** 10.1101/2023.07.19.549235

**Authors:** Julien Castel, Guangping Li, Oriane Onimus, Emma Leishman, Patrice D. Cani, Heather Bradshaw, Ken Mackie, Amandine Everard, Serge Luquet, Giuseppe Gangarossa

## Abstract

The *N*-acyl phosphatidylethanolamine-specific phospholipase D (NAPE-PLD) catalyzes the production of *N*-acylethanolamines (NAEs), a family of endogenous bioactive lipids, which are involved in various biological processes ranging from neuronal functions to energy homeostasis and feeding behaviors. Reward-dependent behaviors depend on the dopamine (DA) transmission between the ventral tegmental area (VTA) and the nucleus accumbens (NAc) which conveys reward-values and scales reinforced behaviors. However, whether and how NAPE-PLD may contribute to the regulation of feeding and reward-dependent behaviors has not been investigated yet. This biological question is of paramount importance since NAEs are altered in obesity and metabolic disorders.

Here, we show that transcriptomic meta-analysis highlights a potential role for NAPE-PLD within the VTA→NAc circuit. Using brain-specific invalidation approaches, we report that the integrity of NAPE-PLD is required for the proper homeostasis of NAEs within the midbrain VTA and it affects food-reward behaviors. Moreover, region-specific knock-down of NAPE-PLD in the VTA resulted in enhanced food-reward seeking and reinforced behaviors which were associated with increased *in vivo* DA release dynamics in response to both food and non-food-related rewards together with heightened tropism towards food consumption. Furthermore, midbrain knock-down of NAPE-PLD, which led to increased energy expenditure and adapted nutrients partitioning, elicited a relative protection against high-fat diet-mediated body fat gain and obesity-associated metabolic features.

In conclusion, these findings unravel a new key role of VTA NAPE-PLD in shaping DA-dependent events, feeding behaviors and energy homeostasis, thus providing new insights on the regulation of body metabolism.

**Highlights:**

- NAPE-PLD and NAEs are enriched in the VTA and regulate food-reinforced behaviors and reward processes.
- NAPE-PLD scales *in vivo* VTA→NAc dopamine dynamics.
- NAPE-PLD in the VTA contributes to whole-body energy balance and metabolic efficiency.
- Downregulation of VTA NAPE-PLD ameliorates obesity-associated metabolic features.

## Introduction

The regulation of feeding behaviors and energy homeostasis is a cardinal and evolutionarily conserved physiological feature in mammals. By mobilizing several and functionally distinct brain circuits (1,2), such regulation tightly depends on metabolic and nutritional demands as well as on the reinforcing and hedonic properties of foods. Among the different regulatory pathways which signal homeostatic states and scale feeding behaviors, lipids, either nutritional and/or endogenous species, represent key mediators in gating the functional adaptability of complex neuronal networks that synchronize food intake and energy expenditure (3,4).

Endogenous bioactive lipids are a major class of biologically active mediators with a critical role in the regulation of several functions spanning from homeostasis to cognition. Among these, the membrane phospholipid-derived long-chain fatty acids *N*-acylethanolamines (NAEs) are potent signaling molecules in peripheral tissues as well as in the central nervous system (CNS). Given their wide distribution and multiple biological functions, the modulation of NAEs tone represents an interesting target for the development of new therapeutic approaches (5).

Among NAEs, the *N*-arachidonoylethanolamine (AEA or anandamide), a *bona fide* endocannabinoid (eCB) agonist of the cannabinoid receptors CB1R and CB2R, is synthetized by the Ca^2+^-dependent enzyme *N*-acyl phosphatidylethanolamine-specific phospholipase D (NAPE-PLD) (6,7) from the membrane *N*-acylphosphatidylethanolamine (NAPE). However, the enzyme NAPE-PLD is also important for the synthesis of other NAEs that do not bind to CBRs, including *N*-oleoylethanolamine (OEA), *N*-palmitoylethanolamine (PEA) and *N*-stearoylethanolamine (SEA) (5). In fact, while AEA binds to the cannabinoid receptors CB1R, CB2R and the transient receptor potential vanilloid 1 (TRPV1) (8), OEA, PEA and SEA activate several non-cannabinoid receptors, including peroxisome proliferator-activated receptors (PPARα), GPR55 and GPR119 (9,10). Importantly, these NAEs are involved in the regulation of appetite and energy homeostasis through different mechanisms (11–15), notably by acting within the gastrointestinal tract (13,16), at the gut-brain interface (9,17,18) and/or directly in the brain (19–21). While NAPE-PLD expression, enzymatic activity and related byproducts have been detected in the mouse, rat and human brains (7,22–24), the key functions of this enzyme within the brain remain still elusive and not well-elucidated. In fact, most of the current literature has focused on distinct NAEs based on their transducing effectors (receptors, transcription factors) and not on the enzyme itself. Only recently, a few studies have shown that pharmacological blockade of NAPE-PLD activity (25) or selective ablation of NAPE-PLD in stress-activated neurons (21) impaired limbic functions (fear extinction, anxiety) mainly through the regulation of the hypothalamus-pituitary-adrenal (HPA) axis, thus highlighting the importance of NAPE-PLD and its NAEs in brain functions. Furthermore, NAPE-PLD silencing and consequent increased availability of its NAPE substrate are neuroprotective in response to 6-OHDA-induced loss of dopamine (DA)-neurons (20), suggesting an important role of NAPE-PLD in regulating physiological and cellular functions of DA-neurons.

Moreover, the developmental invalidation of the Napepld gene alters very-long chain fatty acids composition in the brain suggesting a more complex role for NAPE-PLD than previously appreciated and the existence of alternative biosynthetic pathways (26,27). Furthermore, the non-CB1R-related signaling engaged by fatty acid ethanolamines (FAEs) produced by the specific enzymatic activity encoded by the Napepld gene (ID: 242864) points toward a role for this enzyme in the generation of bioactive FAEs with a wide array of functions including the regulation of energy balance (OEA), inflammation (PEA) or pain sensitivity (SEA).

Interestingly, the presence of a single-nucleotide polymorphism (SNP) on the coding region of the Napepld gene (rs17605251) has been associated with severe obesity (BMI ≥ 35 kg/m^2^) (28,29), suggesting a potential role for NAPE-PLD in the regulation of energy homeostasis, food-motivated behaviors and metabolic disorders.

In the present study, we took advantage of several integrative *in vivo* approaches following genetic and/or viral induction of tissues-specific deletion of NAPE-PLD in basal, food-motivated and obese conditions. Within the mouse mesolimbic reward system, notably the ventral tegmental area (VTA), we found that the enzyme NAPE-PLD functions as a fine-tuning gatekeeper of reward events and dopamine dynamics as well as an important regulator of energy homeostasis and metabolic efficiency in both physiological and pathological (obesity) contexts.

## Materials and methods

### Animals

All experimental procedures were approved by the Animal Care Committee of the Université Paris Cité (CEB-06-2017, APAFiS #11033) and carried out following the 2010/63/EU directive. 8-20 weeks old transgenic male or female mice were used and housed in a room maintained at 22 +/-1 °C, with a light period from 7h00 to 19h00. Regular chow diet (3.24 kcal/g, reference SAFE® A04, Augy, France) and water were provided ad libitum unless otherwise stated. Diet-induced obesity (DIO) was achieved by exposing the mice to a 3-4 months period of high-fat diet (HFD, #D12492, Research Diets Inc., 5.24 kcal/g). The following transgenic mouse lines were used: Napepld^f/f^ (18,30), Nes-Cre (31,32) [JAX003771, B6.Cg-Tg(Nes-cre)1Kln/J], Villin-Cre^ERT2^ (33) [JAX020282, B6.Cg-Tg(Vil1-cre/ERT2)23Syr/J] and Pgk-Cre (34) [JAX020811, B6.C-Tg(Pgk1-cre)1Lni/CrsJ]. All mouse lines were backcrossed and maintained as C57Bl6/J. Breeding-derived lines were generated and genotyped in house.

### Drugs

GBR12909 (10 mg/kg, Tocris) and cocaine hydrochloride (15 mg/kg, Sigma-Aldrich) were dissolved in saline. Tamoxifen (MP Biomedicals) was suspended in ethanol (100 mg/ml, stock solution). A ready-to-use 10 mg/ml tamoxifen solution was prepared by adding filtered sunflower oil, followed by 30 min sonication and stored at 4 °C for up to 1 week. Tamoxifen solution was sonicated 5 min before administration and injected for 5 consecutive days (100 µl/day, i.p.).

### Viral constructs

pAAV-hsyn-GRAB_DA2m was a gift from Yulong Li (Addgene plasmid #140553; http://n2t.net/addgene:140553; RRID:Addgene_140553).

pAAV.CMV.HI.eGFP-Cre.WPRE.SV40 was a gift from James M. Wilson (Addgene plasmid #105545; http://n2t.net/addgene:105545; RRID:Addgene_105545).

pAAV.CMV.PI.eGFP.WPRE.bGH was a gift from James M. Wilson (Addgene plasmid #105530; http://n2t.net/addgene:105530; RRID:Addgene_105530)

### Stereotaxic surgery

For all surgical procedures, mice were rapidly anesthetized with isoflurane (3%, induction), injected (ip) with the analgesic buprenorphine (Buprecare, 0.3 mg/kg, Recipharm, Lancashire, UK) and ketoprofen (Ketofen, 10 mg/kg, France), and maintained under isoflurane anesthesia (1.5%) throughout the surgery.

#### Stereotaxic surgery

Mice were placed on a stereotactic frame (David Kopf Instruments, California, USA). Bilateral (AAV-GFP or AAV-Cre-GFP in the VTA, 0.2 µl/side) or unilateral (GRAB-DA2m in the NAc, 0.3 µl) micro-injections were performed at the following coordinates (in mm from bregma): VTA (L= +/-0.45; AP= -3.4, V= -4.3) and NAc (L= -0.9; AP= +1.18, V= -4.4). All viruses were injected at the rate of 0.05 µl/min. Mice recovered for at least 3-4 weeks after the surgery before being involved in experimental procedures.

### *In vivo* fiber photometry

For *in vivo* dopamine imaging (GRAB-DA2m, (35)), a chronically implantable cannula (Doric Lenses, Québec, Canada) composed of a bare optical fiber (400 µm core, 0.48 N.A.) and a fiber ferrule was implanted 100 µm above the location of the viral injection site in the NAc. The fiber was fixed onto the skull using dental cement (Super-Bond C&B, Sun Medical). Real time fluorescence was recorded using fiber photometry as described in (36,37). Fluorescence was collected in the NAc using a single optical fiber for both delivery of excitation light streams and collection of emitted fluorescence. The fiber photometry setup used 2 light emitting LEDs: 405 nm LED sinusoidally modulated at 330 Hz and a 465 nm LED sinusoidally modulated at 533 Hz (Doric Lenses) merged in a FMC4 MiniCube (Doric Lenses) that combines the 2 wavelengths excitation light streams and separate them from the emission light. The MiniCube was connected to a fiber optic rotary joint (Doric Lenses) connected to the cannula. A RZ5P lock-in digital processor controlled by the Synapse software (Tucker-Davis Technologies, TDT, USA), commanded the voltage signal sent to the emitting LEDs via the LED driver (Doric Lenses). The light power before entering the implanted cannula was measured with a power meter (PM100USB, Thorlabs) before the beginning of each recording session. The light intensity to capture fluorescence emitted by 465 nm excitation was between 25-40 µW, for the 405 nm excitation this was between 10-20 µW at the tip of the fiber. The fluorescence emitted by the GRAB was collected by a femtowatt photoreceiver module (Doric Lenses) through the same fiber patch cord. The signal was then received by the RZ5P processor (TDT). On-line real time demodulation of the fluorescence due to the 405 nm and 465 nm excitations was performed by the Synapse software (TDT). Signals were exported to Python 3.0 and analyzed offline as previously described (36). Data are presented as z-score of ΔF/F.

### Metabolic efficiency analysis

Metabolic efficiency was measured as previously described (36). Briefly, mice were monitored for whole energy expenditure (EE), O_2_ consumption, CO_2_ production, respiratory exchange rate (RER=VCO_2_/VO_2_, V=volume), and locomotor activity using calorimetric cages (Labmaster, TSE Systems GmbH, Bad Homburg, Germany). Gases ratio was determined through an indirect open circuit calorimeter. This system monitors O_2_ and CO_2_ at the inlet ports of a tide cage through which a known flow of air is ventilated (0.4 L/min) and regularly compared to an empty reference cage. O_2_ and CO_2_ were recorded every 15 min during the entire experiment. EE was calculated using the Weir equation for respiratory gas exchange measurements. Food intake was measured with sensitive sensors for automated online measurements. Calorimetric studies to investigate voluntary exercise-induced metabolic adaptions were performed in metabolic cages equipped with running wheels (Promethion, Sable Systems, Nevada, USA). Mice were monitored for body weight and composition at the entry and exit of the experiments using an EchoMRI (Whole Body Composition Analyzers, EchoMRI, Houston, USA). Data analysis was performed on Excel XP using extracted raw values of VO_2_ (ml/h), VCO_2_ (ml/h), and EE (kcal/h).

### Oral glucose tolerance test (OGTT)

Animals were fasted 6 hours before oral gavage of glucose (2 g/kg). Blood glucose was measured from the vein blood tail using a glucometer (Menarini Diagnotics, Rungis, France) at 0, 5, 10, 15, 30, 45, 60, 90, and 120 min. Blood samples were taken at 0, 15, 30 and 60 min to measure insulin levels (mouse ultrasensitive insulin ELISA kit, ALPCO, Salem, USA).

### Behaviors

#### Operant conditioning

Mice were food-restricted and maintained at 90% of their initial body weight to facilitate learning and performance during the whole operant conditioning. Computer-controlled operant conditioning was conducted in 12 identical conditioning chambers (Phenomaster, TSE Systems GmbH, Bad Homburg, Germany) during the light phase, at the same hour every day until the end of the procedure. Each operant wall had two levers (one active and one inactive) located 3 cm lateral to a central pellet dispenser. The reinforcer was a single 20-mg peanut butter flavored sucrose tablet (TestDiet, Richmond, USA). Operant training was carried out daily with no interruption for 1h under a fixed-ratio 1 (FR1, 1 lever press=1 pellet). When the discrimination score between active and inactive lever press (active lever presses/inactive lever presses) exceeded chance level, mice were shifted to sessions under a FR5 (5 lever presses=1 pellet) and/or a progressive ratio (PR) [3 lever presses more for each subsequent reinforcer (r=3N+3; N=reinforcer number)]. Whenever of interest, PR was conducted in both food-restricted and sated mice.

#### Conditioned-place preference (CPP)

The CPP paradigm was performed during the light phase either in food-restricted (maintenance at 90% of initial body weight) or normally fed mice. All the compartments were cleaned before each conditioning session. Locomotor activity was recorded with an infrared beam-based activity monitoring system and analyzed with the provided software (Phenomaster, TSE Systems GmbH, Bad Homburg, Germany). The least preferred compartment during the exploration phase was designated as the reward (HFD)-baited compartment whereas the more preferred compartment as the chow-baited compartment (biased protocol). Animals with more than 65% of preference for a compartment on the pre-test day were removed. To reduce anxiety, during the first two days, animals were carefully put in the middle of the apparatus and allowed to freely explore the two compartments for 1h. The subsequent days included alternating conditioning sessions of 1h. After 8 days of conditioning [4 sessions in each compartment (chow and HFD)], animals freely explored the two compartments for 30 minutes. The time spent in the reward-paired compartment before *vs* after conditioning was the primary outcome variable (preference score).

#### T-Maze

Mice were food-restricted (90 % of initial body weight) during the whole paradigm and tested for learning and cognitive flexibility in a T-maze apparatus (arm 35-cm length, 25-cm height, 15-cm width) (38). First, they were habituated to the apparatus (15 min of exploration) for two consecutive days. Then, mice underwent a 5-days training protocol with one arm reinforced with a palatable food pellet (HFD, cat #D12492, 5.24 kcal/g). Each mouse was placed at the starting point and allowed to explore the maze by choosing one of the two arms (reinforced and non-reinforced arms). The chosen arm was then blocked for 20 seconds and the mouse replaced again in the starting arm. This process was repeated for 10 sessions per day. At the end of this training period, cognitive flexibility and relearning processes were assessed in a reversed learning task which consisted in exchanging the reinforced with the non-reinforced arm. Again, mice underwent a 5-days training protocol (10 sessions/day).

#### Time-locked wheel running

Mice had access to a running wheel connected to an automatic revolution counter (Intellibio Innovation) during a limited amount of time (30 min per session, one session per day) during 5 consecutive days.

#### Food preference and choice

Mice were tested for food choice and preference by using non-caloric and/or caloric solutions. Notably, they were exposed to graduated small bottles containing either water, sucralose (2 mM), sucrose (10% w/v) or intralipids 20%. During different days of exposure (1h session with free choice between 2 bottles), preference was measured by comparing the consumption of sucralose *vs* water, sucrose *vs* water and lipids *vs* water.

#### GBR-induced locomotor activity

Locomotor activity induced by GBR12909 (10 mg/kg, i.p.) was recorded in an automated online measurement system using an infrared beam-based activity monitoring system (Phenomaster, TSE Systems GmbH, Bad Homburg, Germany).

### Tissue preparation and immunofluorescence

Mice were anaesthetized with pentobarbital (500 mg/kg, i.p., Sanofi-Aventis, France) and transcardially perfused with cold (4°C) PFA 4% for 5 minutes. Sections were processed and confocal imaging acquisitions were performed as previously described (37,39). GFP staining was not antibody-amplified (AAV-Cre-GFP, AAV-GFP and GRAB-DA2m). The following primary antibodies were used: rabbit anti-TH (1:1000, Merck Millipore, #AB152) and rabbit anti-cFos (1:500, Synaptic Systems, #226003). Quantification of cFos-immunopositive cells was performed using the cell counter plugin of ImageJ taking a fixed threshold of fluorescence as standard reference.

### Lipidomics

Tissue extracts and HPLC/MS/MS were performed as previously described (40). In brief, samples were placed in 50 volumes of HPLC-grade methanol then spiked with 500 pmols deuterium-labeled N-arachidonoyl glycine (d8NAGly; Cayman Chemical, Ann Arbor, MI) as an internal standard to determine extraction efficiency. Samples were placed on ice in darkness for 2 hours then individually homogenized. Homogenates were then centrifuged at 19,000g for 20 minutes at 20°C. Supernatants were decanted and diluted with HPLC H_2_O to make a 75:25 water to supernatant solution. Partial purification was achieved using C-18 solid phase extraction columns (Agilent Technologies, Lake Forest, CA). A series of 4 elutions with 1.5 mL of 60%, 75%, 85%, and 100% methanol were collected for analysis. Samples were analyzed using an Applied Biosystems API 3000 triple quadrupole mass spectrometer with electrospray ionization. 20µL from each elution were chromatographed using an XDB-C18 reversed phase HPLC analytical column (Agilent) and optimized mobile phase gradients.

### Reclustering and transcriptomics meta-analysis

Publicly available transcriptomic data were downloaded from Gene Expression Omnibus (https://www.ncbi.nlm.nih.gov/geo/, GSE137763, GSE168156, GSE64526, GSE114918) and analyzed using a Python 3.0 pipeline generated in line with the original publications.

### Statistics

All data are presented as mean ± SEM. Statistical tests were performed with Prism 7 (GraphPad Software, La Jolla, CA, USA). Detailed statistical analyses are listed in the **Suppl. Table 1**. Normality was assessed by the D’Agostino-Pearson test. Depending on the experimental design, data were analyzed using either Student’s t-test (paired or unpaired) with equal variances, one-way ANOVA or two-way ANOVA. The significance threshold was automatically set at p<0.05. ANOVA analyses were followed by Bonferroni post hoc test for specific comparisons only when overall ANOVA revealed a significant difference (at least p<0.05).

## Results

### NAPE-PLD is functionally expressed in the brain and mediates motivational food-responses

In order to explore the role of the NAEs-synthetizing enzyme NAPE-PLD in food-motivated behaviors, we first addressed the consequence of whole-body NAPE-PLD developmental knock-out in a food-reward seeking behavioral paradigm. Here, we used the Napepld^f/f^ mouse line in which the two LoxP sites span the exon 3 (24,30,41), the gene sequence that encodes for the catalytic activity of the enzyme and that efficiently leads to a reduction of NAPE-PLD-derived bioproducts (18,24,30,41). Napepld^f/f^ mice were bred with mice expressing Cre under the pan-promoter phosphoglycerate kinase 1 (Pgk-Cre) which result in whole-body full knock-out (KO, (34)) of NAPE-PLD in subsequent generations. To study the reinforcing and motivational properties of food, we performed an operant conditioning paradigm where animals were trained, under different schedules, to press a lever to obtain a palatable sugar pellet. In both a fixed ratio 1 schedule (FR1, 1 lever press for 1 sugar pellet during 4 daily sessions) or a progressive ratio schedule (PR), which assesses the motivational component of reinforcement behaviors, Napepld^+/+^ (controls) and Napepld^KO^ mice displayed similar performances (**Suppl. Fig. 1A, B**). These results suggest that either physiological compensations occurred during development, as reported by previous genetic invalidations (27), and/or that, despite the expression of NAPE-PLD in the brain (23), brain NAPE-PLD plays a marginal role in reward-seeking behavior. To disentangle these two hypotheses, we moved to brain-restricted ablation of NAPE-PLD. Ablation of NAPE-PLD in the central nervous system (CNS) was achieved by crossing Napepld^f/f^ mice with mice expressing Cre under the control of the promoter Nestin (Nes^Cre+/-^ mice) (31,32). We observed that CNS genetic deletion of NAPE-PLD was associated to an enhanced response to operant behavior. In fact, under FR1 schedule, Napepld^ΔCNS^ mice (males and females) collected a higher number of pellets and had a higher number of active lever presses (**Fig. 1A, A^1^**). However, this enhanced reward-like phenotype was not related to differences in learning (% of active lever over inactive lever) as both genotypes were characterized by very similar discrimination scores (**Fig. 1A^2^**). Once the operant conditioning established, mice were moved to the PR schedule. Again, we noticed that, despite similar learning scores, Napepld^ΔCNS^ mice showed enhanced performances (number of rewards and active lever presses) compared to control mice (**Fig. 1B**, B^1^, B^2^). To exclude that such phenotype was driven by the presence of Cre (Nes^Cre+/-^ (32,42)) rather than the proper deletion of NAPE-PLD, we performed the same behavioral battery in Nes^Cre-/-^ (controls) and Nes^Cre+/-^ mice which both displayed a very similar phenotype on this paradigm (**Suppl. Fig. 1C, D)**, thus indicating that genetic deletion of neuronal NAPE-PLD is responsible for the enhanced reward-behavior observed in Napepld^ΔCNS^ mice. This result revealed that tissue-specific ablation of NAPE-PLD generates different outcomes than whole-body gene deletion. This finding, which points to an effective role of brain NAPE-PLD in food-reward seeking behaviors, also raises the possibility that the contribution of NAPE-PLD in multiple organs (full KO mice) might lead to physiological adjustments eventually driving opposite consequences on a particular behavioral output with an overall mitigated consequence.

**Figure 1:**
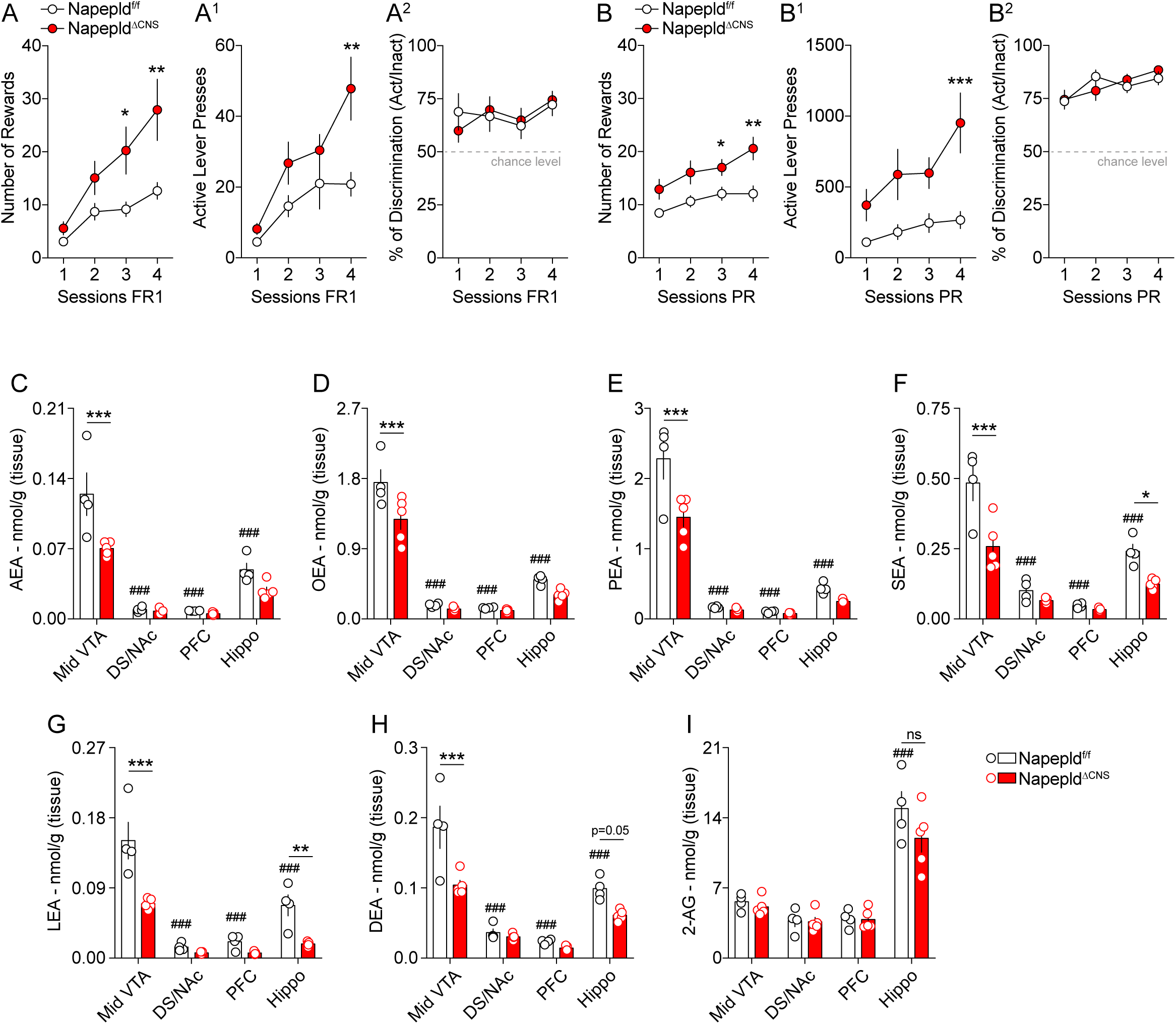
Deletion of NAPE-PLD in the nervous system promotes reward seeking behaviors and reduces *N*-acylethanolamines in the VTA. The reward-like and motivational phenotype of Napepld^f/f^ and Napepld^ΔCNS^ mice was tested through an operant conditioning paradigm (lever press). (A-A^2^) Operant conditioning during 4 consecutive sessions with a fixed ratio 1 (FR1) schedule. (A) Number of rewards, (A^1^) number of active lever presses and (A^2^) percentage of discrimination between the active and inactive lever presses. (B-B^2^) Operant conditioning during 4 consecutive sessions with a progressive ratio (PR) schedule. (B) Number of rewards, (B^1^) number of active lever presses and (B^2^) percentage of discrimination between the active and inactive lever presses. (C-H) Lipidomic analysis of *N*-acylethanolamines (NAEs) and (I) 2-AG in the midbrain ventral tegmental area (VTA), the dorsal striatum/nucleus accumbens (DS/NAc), the prefrontal cortex (PFC) and the hippocampus (Hippo) of Napepld^f/f^ and Napepld^ΔCNS^ mice. Anandamide (AEA), *N*-oleoylethanolamine (OEA), *N*-palmitoylethanolamine (PEA), *N*-stearoylethanolamine (SEA), *N*-linoleoylethanolamine (LEA), *N-*docosahexaenoylethanolamine (DEA), 2-Arachidonoylglycerol (2-AG). Statistics: *p<0.05, **p<0.01 and ***p<0.001 for Napepld^ΔCNS^*vs* Napepld^f/f^ mice; ^###^p<0.001 for VTA *vs* other brain structures (Napepld^f/f^ mice). For number of mice/group and statistical details see **Suppl. Table 1**.

Among the main organs that might contribute to food-dependent reward processes, the gut has emerged as a critical modulator of reinforced behaviors (3,36,43,44). It has been previously shown that mice with a specific and inducible deletion of NAPE-PLD in the intestinal epithelial cells (IEC) (Napepld^ΔIEC^) exhibited a phenotype associated with specific changes in the homeostatic regulation of food intake and altered metabolic adaptations to high-fat diet (18,45). Therefore, we explored the potential contribution of intestinal NAPE-PLD in reward-seeking behavior. Interestingly, Napepld^ΔIEC^ and control mice showed comparable performances in the operant conditioning paradigm (**Suppl. Fig. 1E, F**), indicating that, while intestinal NAPE-PLD is critical for metabolic control (18) and short-term regulation of food intake (45), brain NAPE-PLD might represent a more direct target as acute regulator of food-reward behaviors.

Reinforced behaviors tightly depend on key brain regions that constitute the reward system, notably the midbrain dopamine (DA)-producing ventral tegmental area (VTA) and its dopaminoceptive structures, including the dorsal striatum (DS)/nucleus accumbens (NAc), the prefrontal cortex (PFC) and the hippocampus (Hippo) (46). Therefore, we first investigated whether NAPE-PLD-produced NAEs were altered within these structures in Napepld^ΔCNS^ mice. Lipidomic analyses revealed a significant decrease of several NAEs species [AEA, OEA, PEA, *N*-stearoylethanolamine (SEA), *N*-linoleoylethanolamine (LEA) and *N*-docosahexaenoylethanolamine (DEA)] in the midbrain VTA following CNS deletion of NAPE-PLD (**Fig. 1C-H**). Interestingly, either no major differences (DS/NAc and PFC), specific significant reductions (SEA, LEA and DEA for the hippocampus) or trends of decrease were detected in the levels of NAEs in the other reward-associated brain regions (**Fig. 1C-H**). Of note, no alterations were detected for the endocannabinoid (eCB) 2-AG (**Fig. 1I**). Moreover, within the VTA we did not detect alterations in the levels of other fatty acids (linoleic, arachidonic and oleic acids) (**Suppl. Fig. 2A-C**) or *N*-acylamides (**Suppl. Table 2**), thus indicating that lack of NAPE-PLD specifically affects a subset of endogenous bioactive lipids.

In addition, lipidomic analyses also revealed that NAEs, but not 2-AG (**Fig. 1I**), levels were higher in the VTA compared to the DS/NAc, PFC and hippocampus (**Fig. 1C-H**).

These biochemical results, associated to the enhanced phenotype of Napepld^ΔCNS^ mice in the operant reward-driven behavior (**Fig. 1A, B**), prompted us to investigate the structure- and/or cell type-specific expression of NAPE-PLD in both rodents (rat and mouse) and human brains. First, by taking advantage of single-nucleus RNA transcriptomics (snRNA-seq) in the rat NAc (47) (GSE137763) and VTA (48) (GSE168156), we performed a clustering meta-analyses of transcripts encoding for eCBs- and NAEs-producing enzymes (*Napepld*, *Dagla*, *Daglb*) as well as for eCBs- and NAEs-related transducing effectors (*Cnr1*, *Trpv1*, *Gpr119*, *Ppara*, *Pparg*). Compared to *Dagla* and *Daglb*, we observed that accumbal *Drd1*- and *Drd2*-medium spiny neurons (MSNs) expressed low levels of *Napepld* (6% of 2819 *Drd1*-MSNs and 5% of 1993 *Drd2*-MSNs, respectively) (**Fig. 2A, B**). On the contrary, we detected a higher expression of *Napepld* in VTA-neurons (**Fig. 2C-E**). Since the VTA harbors different neuronal cell types (49), we restricted our meta-analysis to VTA DA-, GABA- and glutamate (Glut)-neurons. In the VTA, we again detected higher levels of *Dagla* and *Daglb* but, interestingly, we observed that *Napepld* was mainly present in VTA DA- and Glut-neurons [12% of 399 DA-neurons (**Fig. 2C**) and 20% of 698 Glut-neurons (**Fig. 2E**), respectively], whereas 9% of GABA-neurons (896 cells) were positive for *Napepld* (**Fig. 2D**). Interestingly, this pattern of *Napepld* expression mirrors the higher levels of NAEs observed in the VTA compared to the DS/NAc (**Fig. 1C**). Moreover, the transcriptomic profiling of eCBs- and NAEs-producing enzymes was in line with the meta-analysis of bulk VTA transcriptomics in the murine (50) (*Slc6a3*-bacTRAP mice, GSE64526, **Fig. 2F**) and human midbrains (51) (GSE114918, **Fig. 2G**).

**Figure 2:**
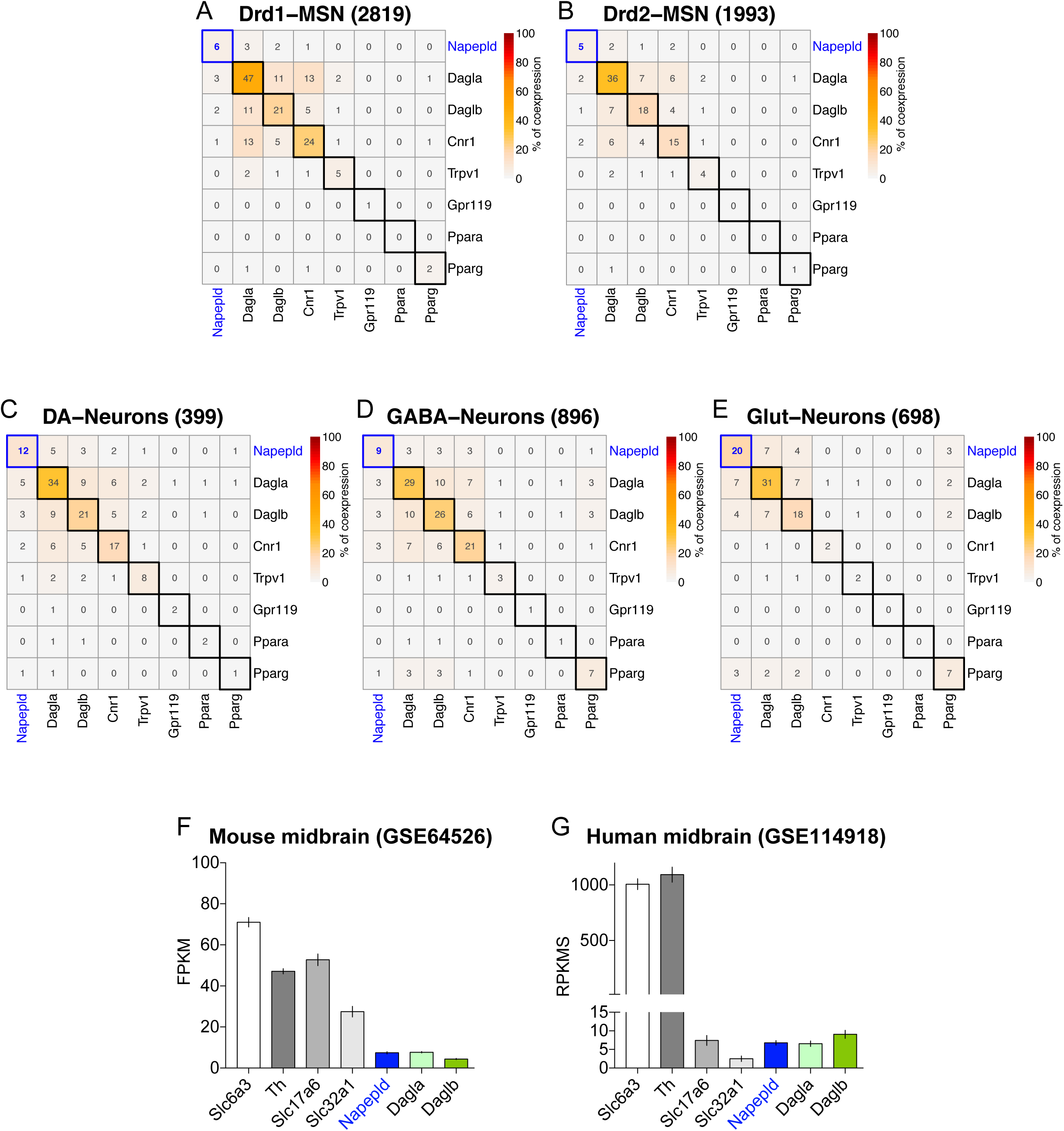
*Napepld* is expressed in the VTA. (A-E) Meta-analysis and clustering of snRNA-seq results in the rat NAc and VTA focusing of NAEs- and endocannabinoids (eCBs)-related synthesis machineries and receptors. Percentage (%) of co-expression of *Napepld* in the NAc *Drd1*-MSNs (A) and *Drd2*-MSNs (B) (extracted from (47)) as well as in the VTA dopamine (DA)-neurons (C), VTA GABA-neurons (D) and VTA glutamate (Glut)-neurons (E) (extracted from (48)). (F) Meta-analysis of bulk transcriptomics in the mouse (extracted from (50)) and (G) human midbrains (extracted from (51)).

Altogether these observations underline the potential role for NAPE-PLD in the midbrain VTA as a regulator of food-associated reward processes.

### VTA NAPE-PLD scales food-motivated behaviors and dopamine releasing dynamics

To precisely interrogate the structure-specific functions of NAPE-PLD in driving food-motivated behaviors, we knocked-down the Napepld gene in the VTA using a local and virally mediated delivery of Cre in the VTA of Napepld^f/f^ mice (**Fig. 3A**). Next, we tested the reinforcing and motivational properties of palatable food using a food-dependent operant conditioning paradigm. In line with the results obtained with Napepld^ΔCNS^ mice (**Fig. 1A, B**), we observed that viral deletion of NAPE-PLD in the VTA promoted food-operant conditioning (increased number of rewards and active lever presses) during both FR1 and PR schedules (**Fig. 3B, C**), with no differences in learning performances as both groups showed similar active/inactive discrimination index (**Fig. 3B, C**).

**Figure 3:**
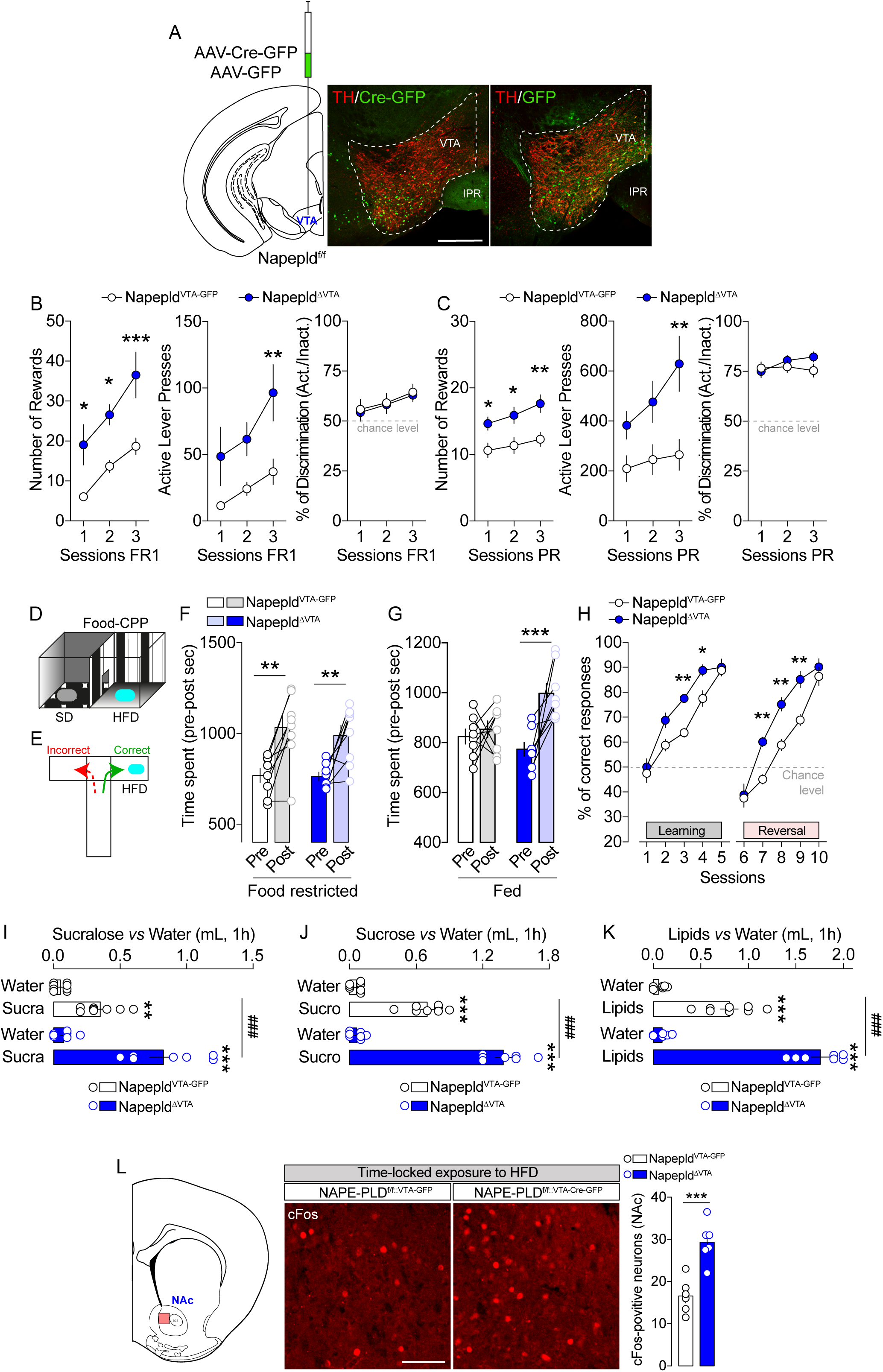
NAPE-PLD knock-down in the VTA promotes reward-seeking behaviors and food preference. (A) Scheme and immunofluorescence sections indicating the viral injection of AAV-Cre-GFP or AAV-GFP in the VTA of Napepld^f/f^ mice. Scale bar: 250 μm. (B, C) Operant conditioning during 3 consecutive sessions with a FR1 (B) and a PR (C) schedule. For both schedules the number of rewards, the number of lever presses and the percentage of discrimination between the active and inactive lever presses are shown. (D, E) Drawings indicate the food-induced conditioned-place preference (CPP) and the T-Maze paradigms, respectively. (F) CPP in Napepld^VTA-GFP^ and Napepld^ΔVTA^ food-restricted mice (10% of body weight reduction). (G) CPP in Napepld^VTA-GFP^ and Napepld^ΔVTA^ sated mice. (H) Learning and reversal learning performances of Napepld^VTA-GFP^ and Napepld^ΔVTA^ food-restricted mice (10% of body weight reduction) in the T-Maze. (I-K) Food preference in Napepld^VTA-GFP^ and Napepld^ΔVTA^ mice using three different choices: sucralose *vs* water (I), sucrose *vs* water (J), and lipids *vs* water (K). (L) Detection and quantification of cFos-positive neurons in the NAc of Napepld^VTA-GFP^ and Napepld^ΔVTA^ mice following the exposure of a fixed amount of high-fat diet (HFD). Scale bar: 100 μm. Statistics: *p<0.05, **p<0.01 and ***p<0.001 for Napepld^ΔVTA^ *vs* Napepld^VTA-GFP^ mice (B, C, H, L). Statistics: **p<0.01 and ***p<0.001 for Napepld^ΔVTA^ (post- *vs* pre-test in CPP) or Napepld^VTA-GFP^ (post- *vs* pre-test in CPP) (F, G). Statistics: **p<0.01 and ***p<0.001 for Napepld^ΔVTA^ (sucra/sucro/lipids *vs* water) or Napepld^VTA-GFP^ mice (sucra/sucro/lipids *vs* water) (I, J, K); ^###^p<0.001 for Napepld^ΔVTA^ *vs* Napepld^VTA-GFP^ mice (I, J, K). For number of mice/group and statistical details see **Suppl. Table 1**.

This enhanced reward phenotype was also present following a FR1→FR5→PR training schedule (**Suppl. Fig. 3A-C**, food restriction) and even in sated conditions (**Suppl. Fig. 3D**), therefore excluding the potentially confounding effect of hunger onto motivational drive. Importantly, this phenotype was also confirmed in female mice (**Suppl. Fig. 3E-H**), again in both food-restricted and sated conditions. Of note, in both males and females, no significant differences in initial body weight and body weight loss (food restriction) were observed between experimental groups (**Suppl. Fig. 4A, B**).

Aside from the motivational component, the liking and learning components of feeding are an integral part of food-reward processes (46). These components can be assessed through behavioral measurements of the positive valence assigned to palatable food in the conditioned-place preference (CPP, **Fig. 3D**) and T-maze (**Fig. 3E**) paradigms, which both rely on the association between reward value and context. In food-restricted conditions, we observed an increased and similar CPP score in both Napepld^VTA-GFP^ and Napepld^ΔVTA^ mice (**Fig. 3F**). However, in sated conditions, only Napepld^ΔVTA^ mice showed an HFD-induced increase in CPP score (**Fig. 3G**), indicating enhanced susceptibility to the reinforcing properties of palatable foods. Using the T-maze paradigm, we next assessed the ability and flexibility of mice to actively learn in discriminating between a rewarded (HFD) and a non-rewarded arm. During the learning phase (first 5 days), we observed that both groups showed a progressive increase in correct responses (%) over training days, with Napepld^ΔVTA^ mice performing significantly better than Napepld^VTA-GFP^ control mice (**Fig. 3H**). Then, mice were tested for their flexibility to relearn the task under a reversal learning schedule (in which the food reinforcer was switched to the previously unreinforced arm of the T-maze). While both groups displayed good performance in learning/flexibility, VTA-specific deletion of NAPE-PLD resulted in a better performance with a more rapid acquisition of the correct entry into the reinforced arm as compared to Napepld^VTA-GFP^ control mice (**Fig. 3H**).

Next, we investigated the role of VTA NAPE-PLD in driving palatable food preference during a time-locked window (1h of exposure). First, we tested the reinforcing properties of the non-caloric sweetener sucralose (2 mM). As shown in **Fig. 3I**, Napepld^ΔVTA^ mice consumed more sucralose than Napepld^VTA-GFP^ control mice. A very similar pattern of enhanced preference was measured with the natural caloric sugar sucrose (10%, **Fig. 3J**) and with emulsified lipids (20%, **Fig. 3K**).

We therefore decided to investigate whether the enhanced reward-like behavior observed in Napepld^ΔVTA^ mice was associated to an increased activity of the nucleus accumbens (NAc), a region highly innervated by VTA projections and whose activity is correlated with food-reward processes (46). However, the enhanced neural response within the reward system might result either from the higher tropism/consumption of palatable food of Napepld^ΔVTA^ mice or from the increased rewarding value despite a fixed amount of food-reinforcer. In order to dissociate these two possibilities, we exposed our experimental groups (sated conditions) to an equal amount of HFD during a time-locked window (1h during which all mice consumed the HFD pellet) and then we analyzed the induction of cFos, a molecular proxy of neuronal activity, in the NAc (**Fig. 3L**). Interestingly, we detected more cFos-positive neurons in the NAc of Napepld^ΔVTA^ mice (**Fig. 3L**), thereby indicating an enhanced responsiveness of the VTA→NAc mesolimbic axis to an equal amount of food-reward consumption.

These results led us to hypothesize that VTA NAPE-PLD and its local NAEs bioproducts may contribute to the regulation of DA dynamics within the VTA→NAc mesolimbic axis. To test this hypothesis, we took advantage of *in vivo* fiber photometry coupled to virally expressed DA biosensors (GRAB-DA2m (35)) to measure DA dynamics in the NAc of Napepld^VTA-GFP^ and Napepld^ΔVTA^ mice (**Fig. 4A, B**). First, we observed that exposing both fasted (**Fig. 4C, D**) and *ad libitum* fed mice (**Fig. 4E, F** and **Suppl. Fig. 5A**) to HFD triggered a higher DA accumulation/release in the NAc of Napepld^ΔVTA^ mice compared to control animals.

**Figure 4:**
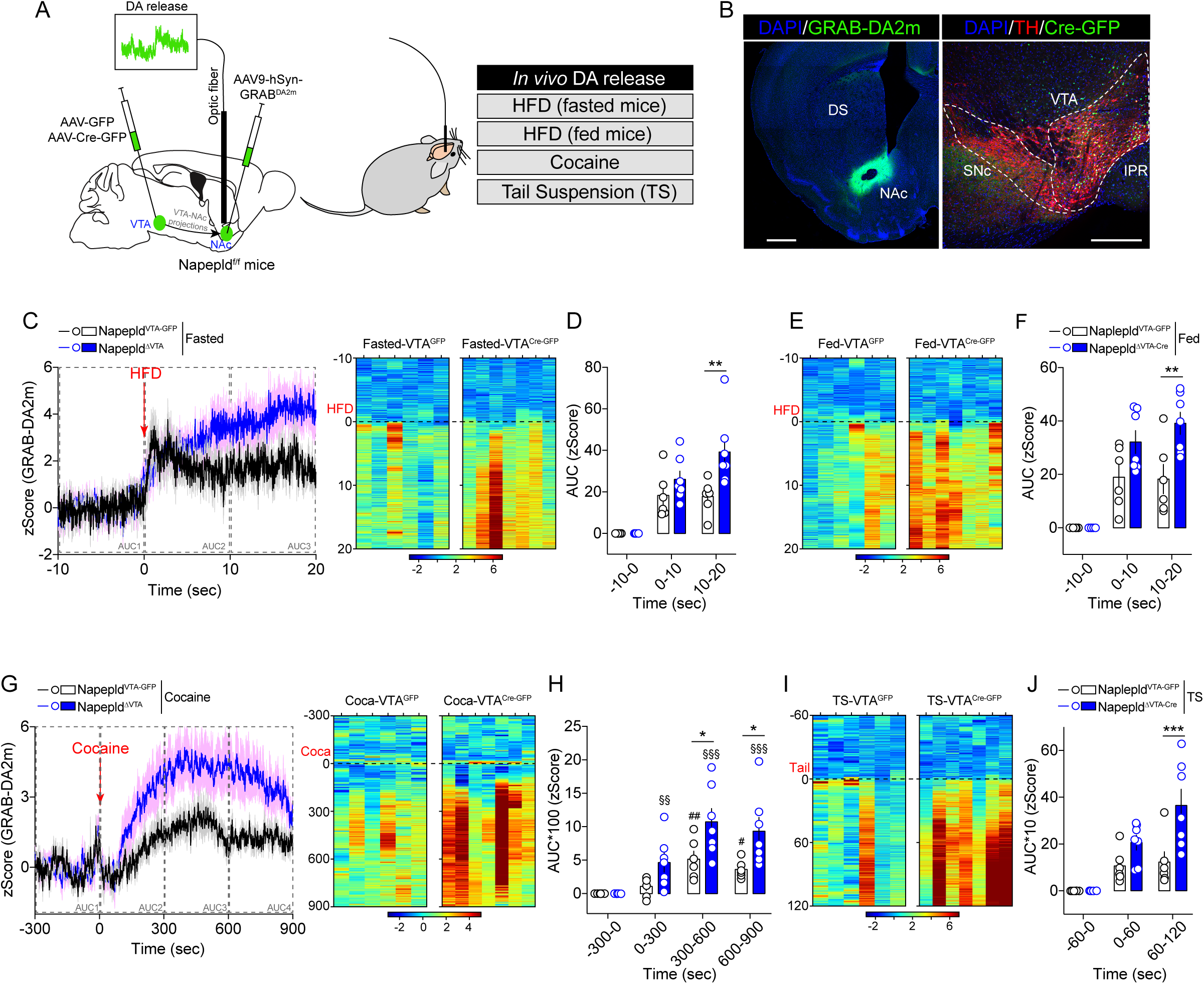
NAPE-PLD knock-down in the VTA regulates *in vivo* DA release dynamics. (A) Drawing indicates the double viral strategies to record *in vivo* DA dynamics in Napepld^VTA-GFP^ and Napepld^ΔVTA^ behaving mice by using fiber photometry coupled to the DA biosensor GRAB-DA2m. DA dynamics were measured during food- and non-food-dependent behaviors. (B) Immunofluorescence detection of GRAB-DA2m in the NAc and AAV-Cre-GFP in the VTA (with TH staining). Scale bars: 250 μm. (C-J) Temporal dynamics and/or heatmaps of DA releasing dynamics in Napepld^VTA-GFP^ and Napepld^ΔVTA^ mice during consumption of HFD in both fasted (C, D) and fed (E, F) conditions as well as during cocaine administration (G, H) and tail suspension (I, J). Statistics: *p<0.05, **p<0.01 and ***p<0.001 for Napepld^ΔVTA^ *vs* Napepld^VTA-GFP^ mice; ^#^p<0.05 and ^##^p<0.01 for cocaine time course in Napepld^VTA-GFP^ mice; ^§§^p<0.01 and ^§§§^p<0.01 for cocaine time course in Napepld^ΔVTA^ mice. For number of mice/group and statistical details see **Suppl. Table 1**.

Second, to further explore whether and how NAPE-PLD may contribute to the regulation of DA-dependent events, we tested *in vivo* DA dynamics also in two non-food-dependent paradigms: the administration of cocaine (**Fig. 4G, H**) and the tail suspension (TS, **Fig. 4I, J** and **Suppl. Fig. 5B**). In both cases, we observed that Napepld^ΔVTA^ mice were characterized by an enhanced accumulation/release of DA in the NAc than Napepld^VTA-GFP^ mice. Lastly, we administered the selective DAT blocker GBR12909 and noticed an enhanced locomotor response in Napepld^ΔVTA^ mice compared to controls (**Suppl. Fig. 5C**), further confirming an amplified DA release/tone as a consequence of VTA NAPE-PLD knock-down.

Overall, these results indicate that VTA NAPE-PLD tightly contributes in orchestrating the responses of midbrain DA-neurons to both food- and non-food-related reinforcers by promoting and boosting the release of DA at VTA→NAc synapses.

### VTA NAPE-PLD contributes to the regulation of food intake and energy homeostasis

Although the regulation of energy homeostasis has been classically ascribed to the hypothalamus and the brainstem (1), new evidence indicates that the reward system also strongly contributes in scaling whole-body metabolic functions (38,52). We therefore explored the metabolic consequences of VTA NAPE-PLD knock-down in the regulation of whole-body metabolic efficiency and peripheral substrates utilization by using longitudinal measurements of indirect calorimetry. As previously observed (**Suppl. Fig. 4**), no major differences were observed in body weight and body composition between the two experimental groups (**Fig. 5A**). However, Napepld^ΔVTA^ mice displayed a spontaneous increase in locomotor activity and in cumulative food intake compared to Napepld^VTA-GFP^ mice during both the light and dark circadian phases (**Fig. 5B, C**). These phenotypes were associated with an overall enhanced energy expenditure (**Fig. 5D**) and to a change in peripheral substrates utilization favoring carbohydrates over lipids-based substrates as indicated by the increase in respiratory exchange ratio (RER, 1=glucose substrate, 0.7=lipids substrate) during the light phase (**Fig. 5E**) and the consequent decrease in fatty acid oxidation (FAO) in Napepld^ΔVTA^ mice during both the light and dark phases (**Fig. 5F**). This feature was also associated with enhanced glucose tolerance during an oral glucose tolerance test at the expense of lower insulin release, suggesting enhanced whole-body glucose dynamics and insulin sensitivity (**Suppl. Fig. 6**).

**Figure 5:**
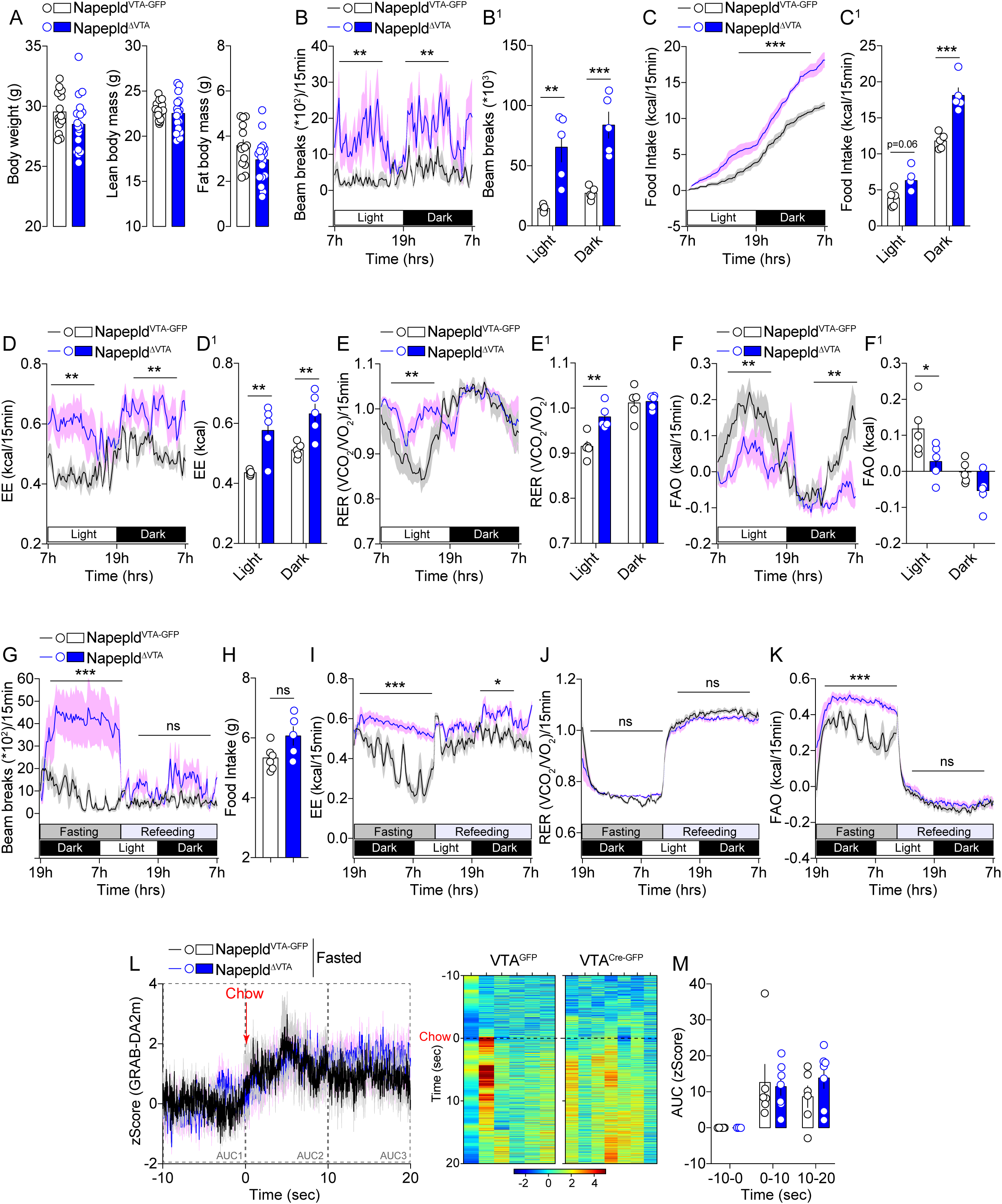
VTA NAPE-PLD contributes to the regulation of energy balance and metabolic efficiency. (A) Body weight and body composition of Napepld^VTA-GFP^ and Napepld^ΔVTA^ mice. (B-K) Indirect calorimetric studies in Napepld^VTA-GFP^ and Napepld^ΔVTA^ mice to assess energy balance and metabolic efficiency. Longitudinal measurements in calorimetric chambers of locomotor activity (B, B^1^), food intake (C, C^1^), energy expenditure EE (D, D^1^), respiratory exchange ratio RER (E, E^1^) and fatty acid oxidation FAO (F, F^1^) in Napepld^VTA-GFP^ and Napepld^ΔVTA^ sated mice. (G-K) Indirect calorimetric studies in Napepld^VTA-GFP^ and Napepld^ΔVTA^ mice undergoing a fasting/refeeding metabolic challenge. (G) Locomotor activity, (H) cumulative food intake, (I) energy expenditure, (J) respiratory exchange ratio and (K) fatty acid oxidation. (L, M) Temporal kinetics and heatmaps of *in vivo* DA release dynamics in Napepld^VTA-GFP^ and Napepld^ΔVTA^ fasted mice and exposed to a chow pellet. Statistics: *p<0.05, **p<0.01 and ***p<0.001 for Napepld^ΔVTA^ *vs* Napepld^VTA-GFP^ mice. For number of mice/group and statistical details see **Suppl. Table 1**.

We then decided to investigate how Napepld^ΔVTA^ mice adapted during manipulation of nutrients availability. We noticed that during a food deprivation period (overnight fasting) Napepld^ΔVTA^ mice were still characterized by increased locomotor activity (**Fig. 5G**) and energy expenditure (**Fig. 5I**), but with no differences in RER (**Fig. 5J**). While the capability to mobilize lipids-based substrates during the fasting-induced lipolysis was similar between the two groups, as indicated by the RER (**Fig. 5J**), the proportion of lipids used a primary source of fuel was enhanced in the fasting period as indicated by the FAO (**Fig. 5K**), thus indicating a metabolic shift toward lipids-based substrates utilization. Interestingly, upon refeeding, mice displayed similar food intake (**Fig. 5H**), locomotor activity (**Fig. 5G**) or substrates utilization (**Fig. 5J, K**), while a slight increase in energy expenditure was still detected in Napepld^ΔVTA^ mice (**Fig. 5I**). These results confirmed the hypothesis that the integrity of NAPE-PLD within the VTA was required for the proper metabolic adaptation to changes in nutrients availability.

Since fasting increases the motivational drive and responsiveness to food, we wondered whether the expression of NAPE-PLD was required to promote DA releasing dynamics in fasted mice exposed to a chow pellet. In contrast to the acute response to palatable HFD (**Fig 4C-F**), consumption of a chow pellet resulted in similar DA releasing dynamics in both Napepld^VTA-GFP^ and Napepld^ΔVTA^ mice (**Fig. 5L, M**). This may suggest that (*i*) the action of VTA NAPE-PLD in the modulation of adaptive metabolic responses to nutritional manipulations can be dissociated from DA release in the fast-refeeding transition and/or (*ii*) VTA NAPE-PLD plays an active role in discriminating between palatable (HFD, **Fig 4C-F**) and regular (chow, **Fig. 5L, M**) foods through the control of reward-dependent DA dynamics.

### VTA NAPE-PLD does not contribute to exercise-motivated behaviors but still regulates energy homeostasis

In mammals, exercise can function as a rewarding/motivational stimulus (53) and the eCBs system, especially within the VTA, has been identified as a key regulator of exercise-induced reinforced behaviors (54–56). We thus decided to extend our investigations to exercise-motivated behaviors (*i*) to investigate whether VTA NAPE-PLD was also important in mediating the reinforcing properties of exercise and (*ii*) to study whether metabolic adaptations observed in Napepld^ΔVTA^ mice (**Fig. 5**) were solely dependent on enhanced locomotor activity.

First, we performed a time-locked access (30 min session/day) to a running wheel. Despite both Napepld^VTA-GFP^ and Napepld^ΔVTA^ mice progressively spent more time wheel-running, we surprisingly noticed a reduced performance in Napepld^ΔVTA^ mice compared to control animals (**Fig. 6A, A^1^**). This led us to investigate whole-body metabolism and metabolic efficiency in calorimetric chambers equipped with running wheels.

**Figure 6:**
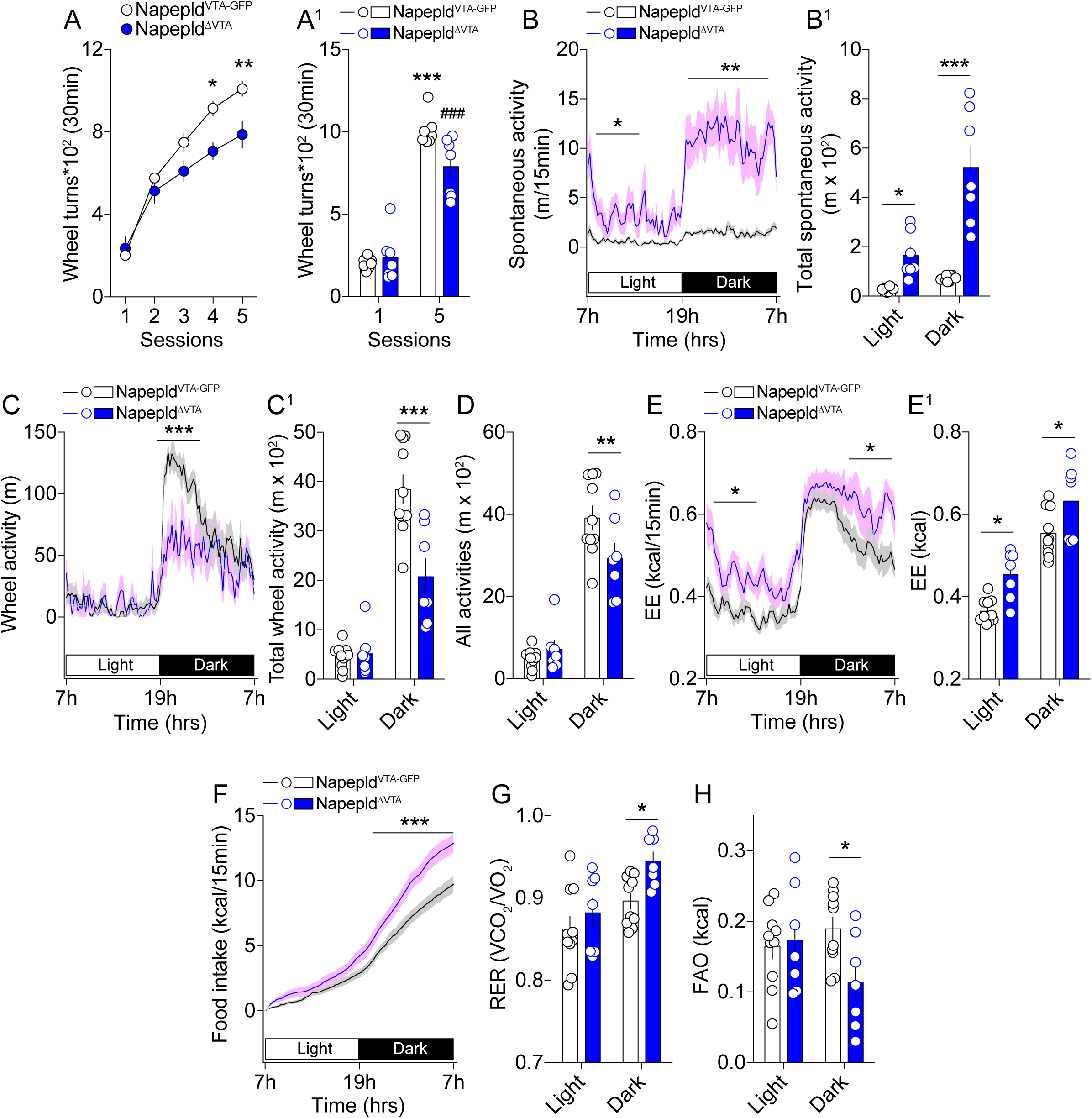
VTA NAPE-PLD regulates energy balance independently from exercise. (A, A^1^) Time-locked (30 min/session) access to running wheels to mimic the reinforcing properties of exercise. (B-H) Indirect calorimetric studies in calorimetric chambers equipped with running wheels to assess energy balance and metabolic efficiency in Napepld^VTA-GFP^ and Napepld^ΔVTA^ mice. (B, B^1^) Spontaneous locomotor activity, (C, C^1^) running wheel activity, (D) total locomotor activity (spontaneous + running wheel activities), (E, E^1^) energy expenditure, (F) cumulative food intake, (G) respiratory exchange ratio and (H) fatty acid oxidation. Statistics: *p<0.05, **p<0.01 and ***p<0.001 for Napepld^ΔVTA^ *vs* Napepld^VTA-GFP^ mice. For number of mice/group and statistical details see **Suppl. Table 1**.

Again, we observed that Napepld^ΔVTA^ mice were characterized by an enhanced spontaneous locomotor activity (**Fig. 6B, B^1^**) and reduced wheel-running activity (**Fig. 6C, C^1^**). When combining both forms of activity (spontaneous + wheel running activities), we detected no differences in the light phase and a lower global activity in Napepld^ΔVTA^ mice during the dark phase (**Fig. 6D**). Of interest, the peculiar metabolic signature associated with VTA NAPE-PLD deletion also remained in this exercise-based paradigm and was characterized by enhanced energy expenditure (**Fig. 6E, E^1^**), food intake (**Fig. 6F**) and RER (**Fig. 6G**), and lower FAO (**Fig. 6H**) in Napepld^ΔVTA^ mice.

Altogether, these results indicate that the role of VTA NAPE-PLD in regulating reward-like processes cannot be generalized to all natural rewards (food *vs* exercise) and that the metabolic adaptations observed in VTA NAPE-PLD-deleted mice are not solely dependent on locomotor activity.

### VTA NAPE-PLD controls metabolic adaptation to obesogenic environment

Food-reward drive, together with changes in metabolic outputs in response to food environment, are important contributors of the obesity pandemics. Given the above-mentioned results showing a key role of VTA NAPE-PLD in controlling reward and metabolic processes, we hypothesized that NAPE-PLD may influence the (mal)adaptive responses to an obesogenic environment. Thus, Napepld^VTA-GFP^ and Napepld^ΔVTA^ mice were chronically exposed to an obesogenic diet (3 months of HFD) and then metabolically characterized.

First, we noticed no significant differences in the body weight and lean mass composition of both HFD-exposed experimental groups (**Fig. 7A**). However, fat body mass was significantly lower in Napepld^ΔVTA^ mice (**Fig. 7A**). The analysis of metabolic efficiency revealed that obese Napepld^ΔVTA^ mice displayed increased nocturnal locomotor activity (**Fig. 7B**) and enhanced nocturnal food intake (**Fig. 7C, C^1^**). Surprisingly, we detected a higher energy expenditure (**Fig. 7D, D^1^**) and FAO (**Fig. 7E, E^1^**) in obese Napepld^ΔVTA^ mice, whereas the RER resulted unchanged (**Fig. 7F**). This metabolic blueprint suggests that, depending on diets (chow *vs* HFD) and metabolic profiles (lean *vs* obese), VTA NAPE-PLD readily allows the plastic adaptation of nutrients partitioning (**Fig. 5E-F** *vs* **Fig. 7E-F**) in order to maintain a higher energy expenditure.

**Figure 7:**
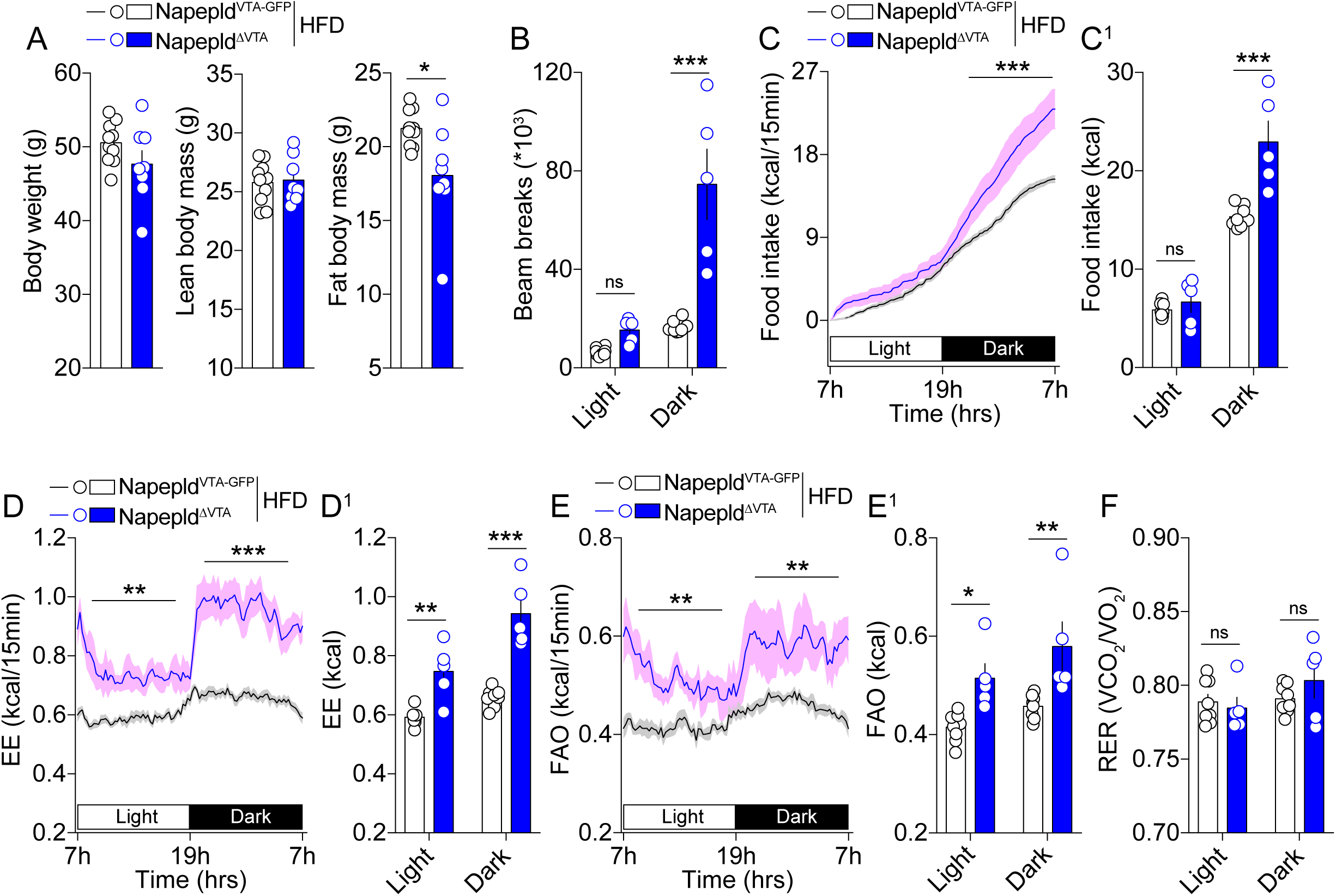
VTA NAPE-PLD protects from obesity-associated metabolic features. (A) Body weight and body composition (fat and lean mass) of Napepld^VTA-GFP^ and Napepld^ΔVTA^ mice exposed to chronic high-fat diet (HFD) and consequently characterized for energy balance and metabolic efficiency. (B) Cumulative locomotor activity, (C, C^1^) food intake, (D, D^1^) energy expenditure, (E, E^1^) fatty acid oxidation and (F) respiratory exchange ratio. Statistics: *p<0.05, **p<0.01 and ***p<0.001 for Napepld^ΔVTA^ *vs* Napepld^VTA-GFP^ mice. For number of mice/group and statistical details see **Suppl. Table 1**.

These results indicate that, within the VTA, the deletion of NAPE-PLD partially protects against diet-induced obesity.

## Discussion

NAEs represent an important family of endogenous bioactive lipids involved in several biological process including adaptive stress responses and emotional states (21,57), pain (58), inflammation (59), feeding and metabolism (9,14,17,18,45). In an effort to characterize the phospholipase D (PLD)-mediated enzymatic activity that converts NAPE into NAEs, *in vitro* studies identified NAPE-PLD as able to produce NAE derivatives from NAPE (7). Genetic invalidation of the Napepld gene (27,30) revealed that, in the brain, several alternative enzymatic pathways exist for the synthesis of polyunsaturated fatty acid NAEs while NAPE-PLD activity seems more critical for saturated/monounsaturated NAEs, with a drastic decrease of these compounds escalating with carbons chain length. Later, in depth lipidomic analysis revealed a broader role for NAPE-PLD with a large spectrum and region-specific consequences in lipidome alteration in the brain (24,30). While constantly evolving, the sensitivity and limitations of quantitative methods to measure NAEs may be in part implicated in the difficulty to formerly assigned a definitive set of substrates and bioproducts to NAPE-PLD. It is now clear that NAPE-PLD bioproducts include important lipid mediators which, by acting on a variety of transcriptional and signaling cascades, control cellular and physiological responses (5).

In the present study we explored the role of NAPE-PLD in food- and reward-driven behaviors. Whole-body deletion of NAPE-PLD did not affect food-reward operant conditioning, suggesting that compensatory mechanisms, as previously reported with a model of developmental NAPE-PLD KO mice (27), may be at play. However, using neural-specific genetic deletion (Nestin-Cre) or midbrain-specific viral invalidation approaches we revealed that the integrity of NAPE-PLD within the midbrain VTA is required for NAEs synthesis (AEA, OEA, PEA, SEA, LEA and DEA) and that NAPE-PLD acts as a gatekeeper for fine-tuning food-reward behaviors and for regulating, at least as a contributor, energy balance and whole-body metabolism. In fact, viral down-regulation of NAPE-PLD in the VTA was associated with an enhanced tropism towards palatable foods and a stronger conditioning for food-related rewards (operant conditioning, conditioned-place preference and T-maze paradigm). These phenomena were associated with an enhanced activity of midbrain VTA DA-neurons and their DA release dynamics in response to food rewards and also to non-food-related stimuli. Consistent with the notion of a region-specific biosynthesis and action of NAEs, RNA-seq meta-analyses revealed low levels of *Napepld* in postsynaptic striatal dopaminoceptive neurons (*Drd1*- and *Drd2*-MSNs), but a higher expression of *Napepld* in VTA-neurons, notably DA- and glutamate-neurons.

Previous studies focusing on nicotine reinforcement and tobacco use disorder (TUD) have shown that pharmacological inhibition of the fatty acid amid hydrolase (FAAH), one of the main enzymes responsible for the degradation of NAEs, reduces nicotine-enhanced DA transmission and nicotine reinforcement (19,60) through the activation of PPARα by OEA/PEA and the activation of intracellular cascades leading to the reduction of nicotinic receptors onto midbrain DA-neurons (19,60). In line with our results showing increased dopaminergic VTA→DA transmission following downregulation of VTA NAPE-PLD, these electrophysiological studies have clearly demonstrated that OEA/PEA inhibit DA-neurons whereas inhibition of PPARα promotes their spontaneous activity (19,61,62). However, these aforementioned reports, although seminal, did not formally test nor established the role of midbrain NAPE-PLD in these processes. In our hands, specific knock-down of NAPE-PLD in the VTA resulted in enhanced DA transmission in response to reward stimuli. In that view, our results are perfectly in line with a putative role for NAPE-PLD-derived substrates as negative modulators of VTA DA-neurons. While our study provides an additional mechanism by pointing at NAPE-PLD has a potential candidate, it also extends the role of this enzyme to the control of DA-dependent behaviors and DA releasing dynamics in response to reward stimuli well-beyond nicotine. Although our results do not rule out whether the NAPE-PLD/NAPE→NAEs machinery controls both the tonic and phasic DA release (63–65), they clearly indicate that VTA NAPE-PLD activity, through the synthesis of NAEs, is an integral component of the control of DA-neurons’ activity. Whether the cellular accumulation of NAPE and/or the decreased levels of NAEs are the primary responsible for the changes in DA-dependent behaviors and DA releasing dynamics is still unknown. Indeed, NAPE-PLD silencing seems to confer a protective action for increased NAPE species in 6-OHDA-induced neural damage (20), suggesting that the regulation of DA-neurons might be linked to NAPE/NAE membrane homeostasis. Given the multiple roles of endogenous bioactive lipids, it is possible that the invalidation of NAPE-PLD in the VTA may lead to an imbalance in the NAPE/NAEs ratio with ultimate consequences on a variety of processes including heightened DA responses to reward stimuli. In addition, NAEs, either cannabinoids-like (AEA) and non-cannabinoids-like (OEA, PEA), may be released and act through several *modus operandi*. In fact, while the eCB AEA is retrogradely released *on demand* (66), the non-eCB NAEs may act anterogradely as suggested by the presence of NAPE-PLD also in pre-synaptic terminals (67), thus potentially modulating both intra-VTA microcircuits and/or VTA-projecting circuits (*i.e.* VTA→NAc). However, the VTA does not only harbor DA-neurons (49,68). Importantly, the relatively high expression of NAPE-PLD in VTA Glut-neurons may also indicate a possible regulatory control of this cell type by the NAPE/NAEs ratio in the observed behavioral and metabolic features. This regulation may be exerted either through the local communication among all VTA-neurons or even through VTA^Glut^→NAc projections. In fact, these glutamatergic projections were recently shown to promote reinforcement independently of DA release (69) and we cannot exclude that NAPE-PLD-dependent mechanisms onto these complex neural networks may result from this additional form of VTA→NAc communication in encoding changes in reward-driven behaviors and metabolic efficiency. Despite this limitation, our *in vivo* imaging results clearly reveal that the VTA→NAc dopaminergic transmission is regulated by the NAPE-PLD. Indeed, further studies are warranted to fully flush out how and to which extent midbrain NAPE-PLD regulates DA events by selectively focusing on the local interconnectivity and interdependency of VTA DA-, Glut- and GABA-neurons.

While our study establishes a direct role for NAPE-PLD in the mesolimbic reward circuit in regulating DA release and DA-dependent behaviors, it also unveils the functional connection between VTA NAPE-PLD activity and the control of whole-body metabolism. NAPE-PLD^ΔVTA^ mice displayed increased spontaneous locomotor activity in both fed and fasting, but not refed, conditions, which is consistent with an enhanced activity of VTA DA-neurons (70). These features were associated with increased cumulative food intake and whole-body energy expenditure and, together with other recent studies (52,71,72), they point to the VTA as an important regulator of energy balance and metabolic efficiency. In fact, on chow diet the overall body weight was only marginally affected in NAPE-PLD^ΔVTA^ compared to control mice, suggesting that increased energy expenditure (EE) was compensated by increased energy intake in a closed and well-balanced homeostatic regulation (**Fig. 5**). During food deprivation, NAPE-PLD^ΔVTA^ mice showed a drastic increase in fasting-induced foraging pointing towards a change in adaptive strategy in response to decreased nutrients availability. This increased activity was associated with increased EE and most likely fueled by enhanced lipids-based metabolism (**Fig. 5**). In mice exposed to HFD, a similar increase in spontaneous activity, EE and FAO was observed together with increased food intake. Since NAPE-PLD^ΔVTA^ mice show increased tropism and responsiveness to HFD and palatable food it is possible that this increase in HFD intake may be the result of enhanced palatability for energy-dense foods. An alternative, and not mutually exclusive, hypothesis would be that increased food intake is a homeostatic mechanism to sustain enhanced EE. Although we do not provide a clear evidence to disentangle these two hypotheses, the fact remains that NAPE-PLD knock-down in the VTA confers a protective phenotype against HFD-induced body fat gain and metabolic alterations (**Fig. 7**). This result nicely echoes a study in humans that has identified a common haplotype of the Napepld gene in severe obesity (28). In face of our results, one should consider that VTA NAPE-PLD deletion led to a protective effect against HFD-mediated fat mass gain and metabolic (mal)adaptations but was accompanied by enhanced reward-driven behaviors. This is important from a translational point of view as obesity is characterized by an alteration of peripheral and central eCBs and NAEs in humans (73–75). Our current results, together with those depicting the role of NAPE-PLD in peripheral tissues, culminate in the elaboration of a complex picture in which organs- and regions-specific homeostasis of NAPE/NAEs underline the complexity by which NAPE-PLD exerts its control onto reward-dependent behaviors, energy balance and body weight control. Indeed, it has been previously shown that mice lacking NAPE-PLD specifically in adipocytes displayed spontaneous obesity, higher fat mass, glucose intolerance, and lower adipocyte browning (41). Moreover, mice lacking NAPE-PLD in the intestinal epithelial cells were more sensitive to HFD-induced body weight gain, fat mass gain and hepatic steatosis (18), a phenomenon partially explained by an alteration in food intake behavior (45).

In conclusion, our study provides a direct evidence for a key role of NAPE-PLD in the control of reward-dependent behaviors, DA dynamics and energy metabolism. The main limitation of this study primarily lies in the lack of a clear cell type-specific identification of cellular and molecular mechanisms occurring within the heterogenous VTA. Given the complexity of NAEs action and the variety of bioactive lipids and signaling cascades associated with NAPE-PLD activity, further research and new investigatory tools will be required to fully apprehend the role of NAPE-PLD and its bioproducts in promoting anti-obesity strategies.

## Acknowledgments

We thank Olja Kacanski for administrative support, Isabelle Le Parco, Aurélie Djemat, Daniel Quintas, Magguy Boa, Ludovic Maingault and Angélique Dauvin for animals’ care, and Florianne Michel for genotyping. We acknowledge the technical platform Functional and Physiological Exploration platform (FPE) of the Université Paris Cité, CNRS, Unité de Biologie Fonctionnelle et Adaptative, the viral production facility of the UMR INSERM 1089 and the animal core facility “Buffon” of the Université Paris Cité/Institut Jacques Monod.

## Author contributions

**Conceptualization**: G.G and S.L.; **Investigation**: J.C., G.L., O.O., A.E., E.L., H.B., K.M., P.D.C., S.L., G.G.; **Formal Data Analysis**: J.C., S.L, G.G., H.B., K.M., A.E.; **Resources:** G.G., S.L., P.D.C.; **Funding acquisition**: G.G., S.L., P.D.C.; **Supervision**: G.G. and S.L.; **Writing – original draft**: G.G. and S.L.; **Writing – review & editing**: all authors.

## Competing interests

P.D.C. and A.E. are inventors on patent applications dealing with the use of specific bacteria and components in the treatment of different diseases. P.D.C. was co-founder of The Akkermansia Company SA and Enterosys. The other authors declare no competing interests.

**Supplementary Figure 1: Tissue-specific deletion of NAPE-PLD and operant behavior.** Operant conditioning for food-seeking (lever press), under both FR1 and PR schedules, was evaluated in different mouse lines. NAPE-PLD was genetically removed from the whole-body (Napepld^KO^ mice, A and B) or from intestinal epithelial cells (Napepld^ΔIEC^, E and F). In addition, operant conditioning was also performed in Nestin-Cre mice (C and D) to discard the potential effect of Cre in the phenotype observed in Napepld^ΔCNS^ mice (**Fig. 1A, B**). For number of mice/group and statistical details see **Suppl. Table 1**.

**Supplementary Figure 2: Levels of fatty acids in the VTA of Napepld**^Δ**CNS**^ **mice.** Lipidomic detection of linoleic (A), arachidonic (B) and oleic (C) acids in the VTA of Napepld^f/f^ and Napepld^ΔCNS^ mice. For number of mice/group and statistical details see **Suppl. Table 1**.

**Supplementary Figure 3: NAPE-PLD knock-down in the VTA promotes reward-seeking behaviors in both males and females.** (A-C) Operant conditioning in Napepld^VTA-GFP^ and Napepld^ΔVTA^ male mice following a FR1→FR5→PR training schedule in food-restricted conditions (reduction of 10% of body weight). Of note, Napepld^VTA-GFP^ and Napepld^ΔVTA^ male mice also underwent a session of PR schedule in sated conditions (D). (E-G) Operant conditioning in Napepld^VTA-GFP^ and Napepld^ΔVTA^ female mice following a FR1→FR5→PR training schedule in food-restricted conditions (reduction of 10% of body weight). Of note, Napepld^VTA-GFP^ and Napepld^ΔVTA^ female mice also underwent a session of PR schedule in sated conditions (H). Statistics: *p<0.05, **p<0.01 and ***p<0.001 for Napepld^ΔVTA^ *vs* Napepld^VTA-GFP^ mice. For number of mice/group and statistical details see **Suppl. Table 1**.

**Supplementary Figure 4: NAPE-PLD knock-down in the VTA does not alter body weight and food restriction-mediated weight loss.** (A) Body weight of Napepld^VTA-GFP^ and Napepld^ΔVTA^ male mice undergoing the operant conditioning paradigm. (B) Body weight of Napepld^VTA-GFP^ and Napepld^ΔVTA^ female mice undergoing the operant conditioning paradigm. Statistics: ***p<0.001 for Napepld^VTA-GFP^ mice (initial *vs* food restriction); ^###^p<0.001 for Napepld^ΔVTA^ mice (initial *vs* food restriction). For number of mice/group and statistical details see **Suppl. Table 1**.

**Supplementary Figure 5: VTA NAPE-PLD and *in vivo* DA dynamics.** (A) Temporal kinetics of *in vivo* DA release in Napepld^VTA-GFP^ and Napepld^ΔVTA^ fed mice exposed to HFD corresponding to **Fig. 4E-F**. (B) Temporal kinetics of *in vivo* DA release in Napepld^VTA-GFP^ and Napepld^ΔVTA^ mice undergoing the tail suspension (TS) test and corresponding to **Fig. 4I-J**. (C) Longitudinal and cumulative measurements of GBR12909-induced locomotor activity in Napepld^VTA-GFP^ and Napepld^ΔVTA^ mice. Statistics: **p<0.01 for Napepld^ΔVTA^ *vs* Napepld^VTA-GFP^ mice (GBR experiment). For number of mice/group and statistical details see **Suppl. Table 1**.

**Supplementary Figure 6: VTA NAPE-PLD contributes to glucose homeostasis.** (A) Blood glucose and (B) insulin levels in Napepld^VTA-GFP^ and Napepld^ΔVTA^ mice after an oral glucose tolerance test (OGTT). Statistics: **p<0.01 for Napepld^ΔVTA^ *vs* Napepld^VTA-GFP^. For number of mice/group and statistical details see **Suppl. Table 1**.

