## Supplementary figures and images for "NAPE-PLD in the ventral tegmental area regulates reward events, feeding and energy homeostasis"

### Suppl. Figure 1

#### Suppl. Figure 1

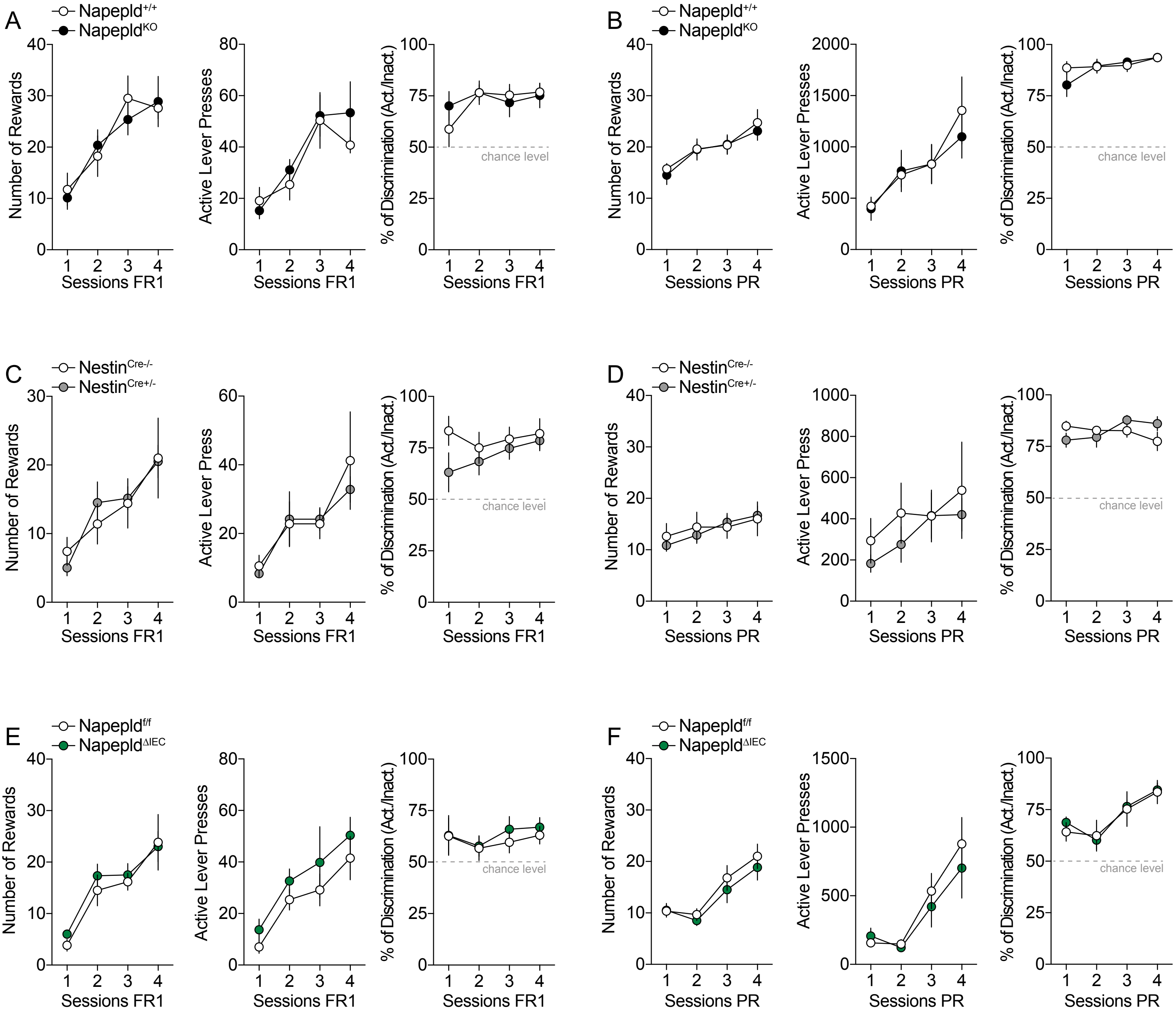

### Suppl. Figure 2

Suppl. Figure 2

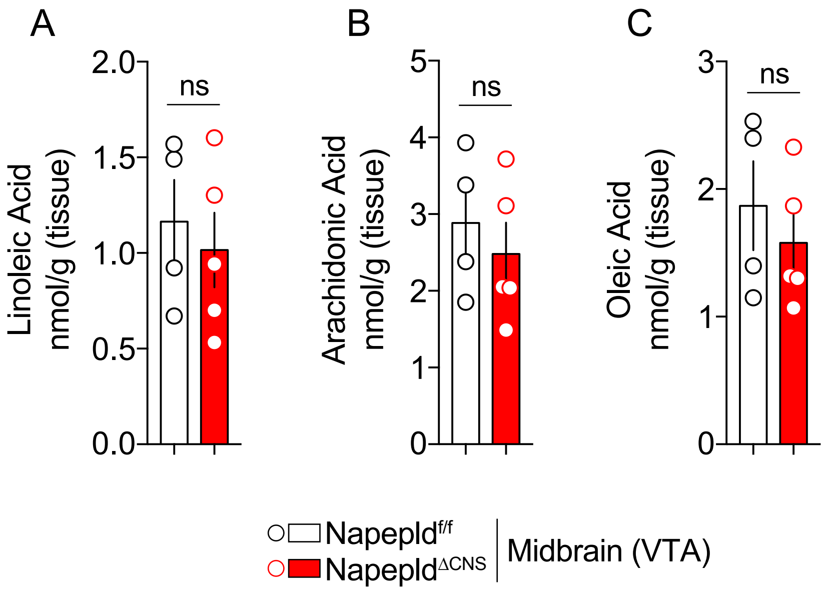

### Suppl. Figure 3

#### Suppl. Figure 3

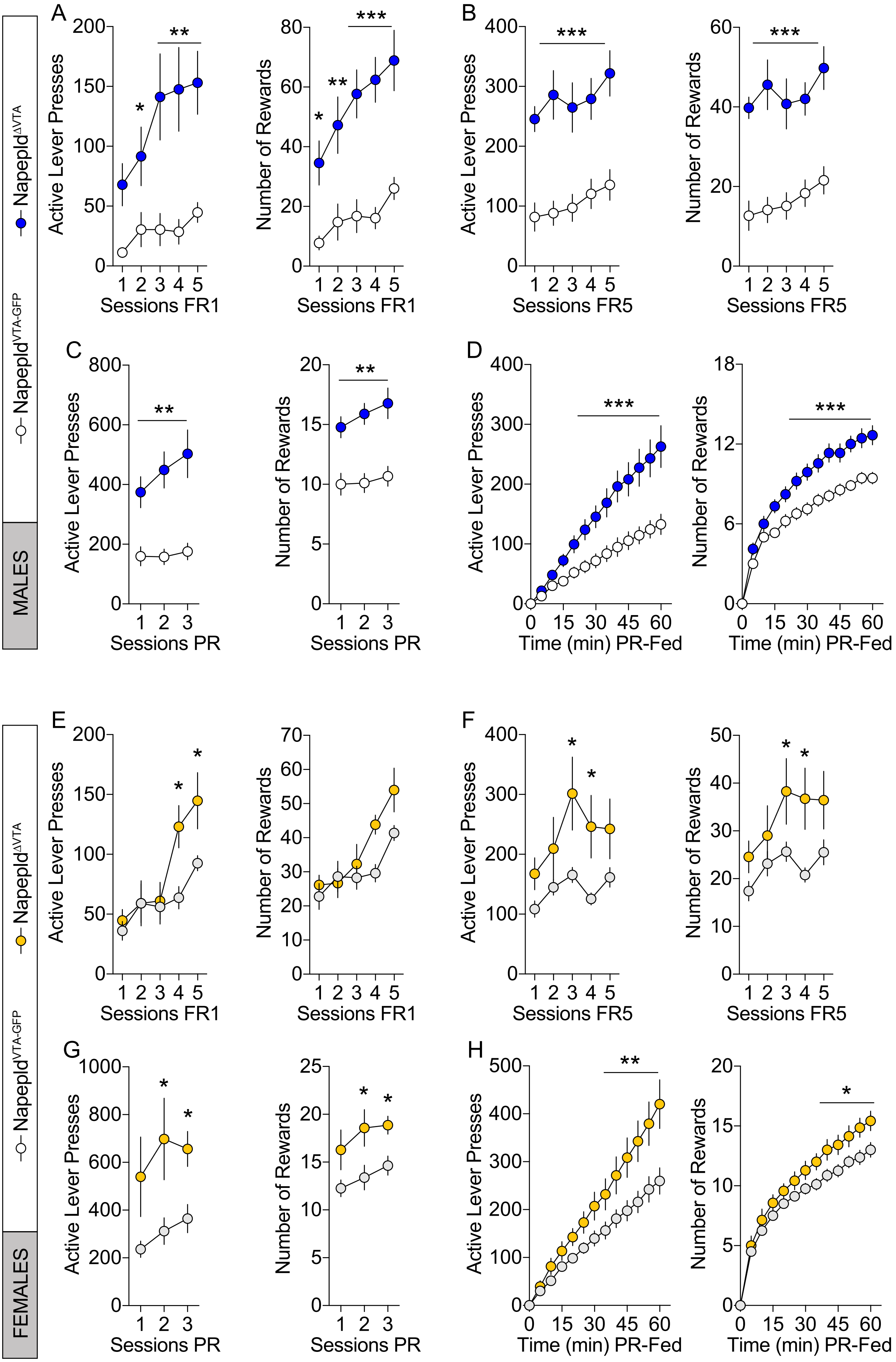

### Suppl. Figure 4

Suppl. Figure 4

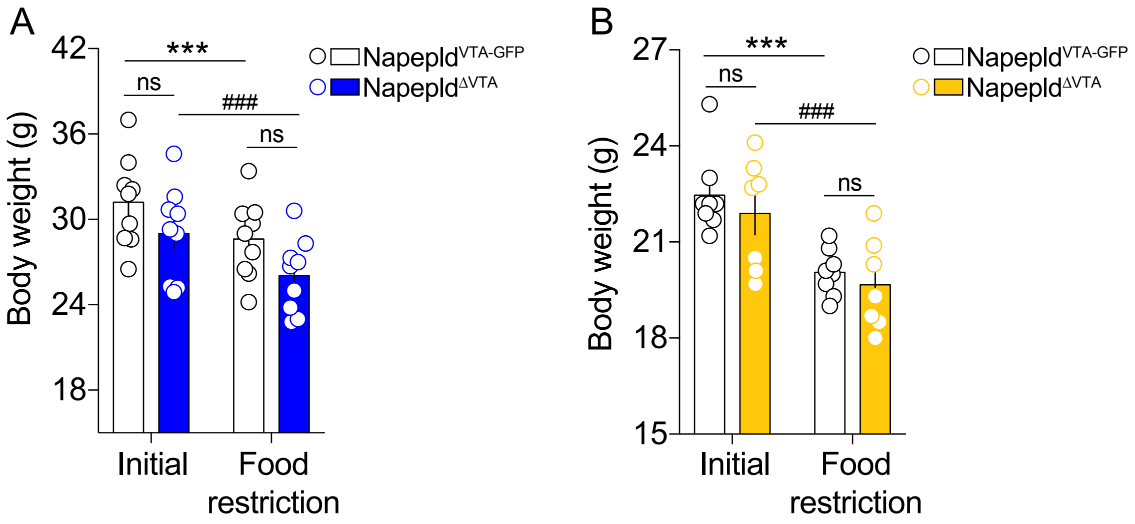

### Suppl. Figure 5

Suppl. Figure 5

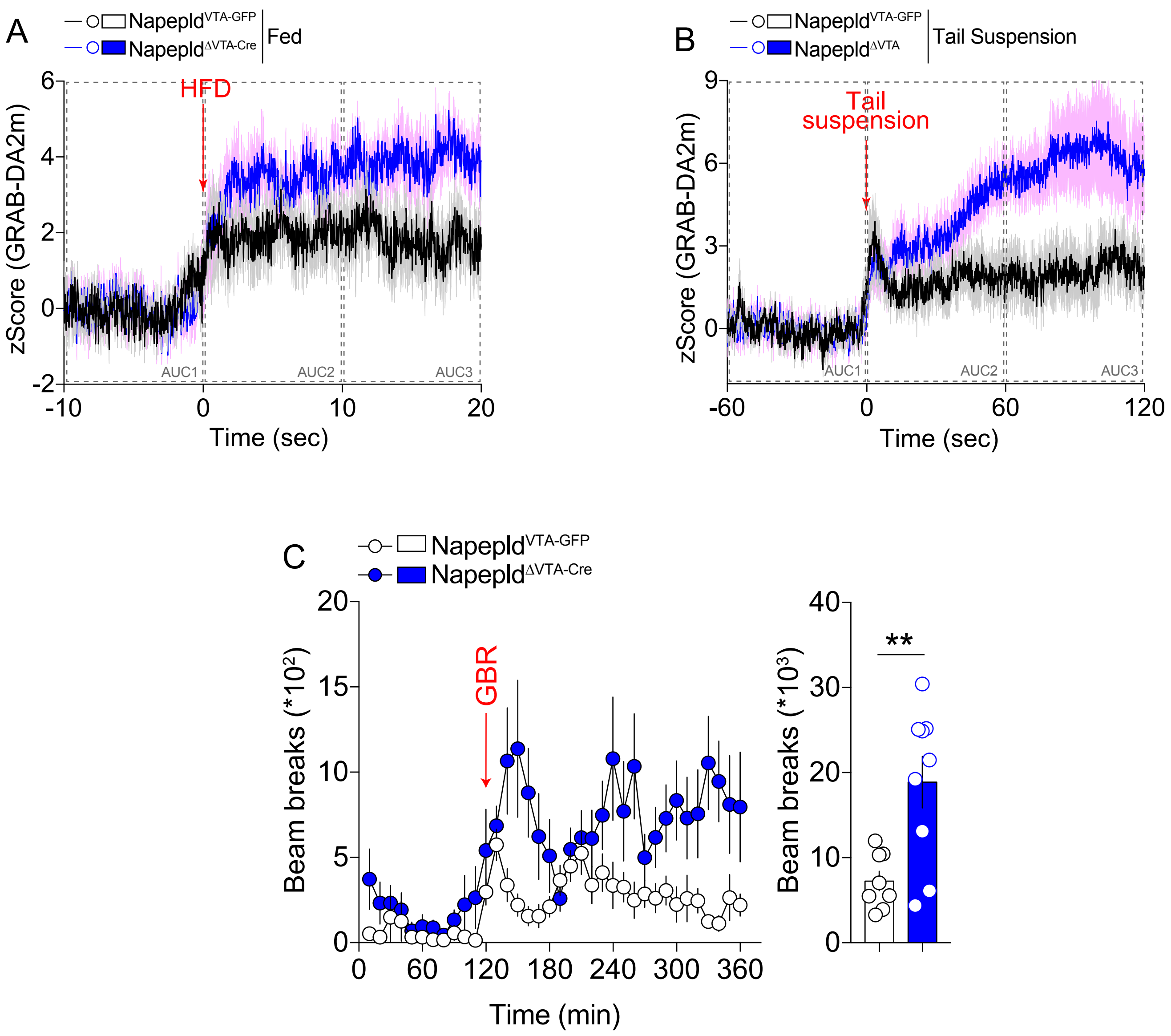

### Suppl. Figure 6

#### Suppl. Figure 6

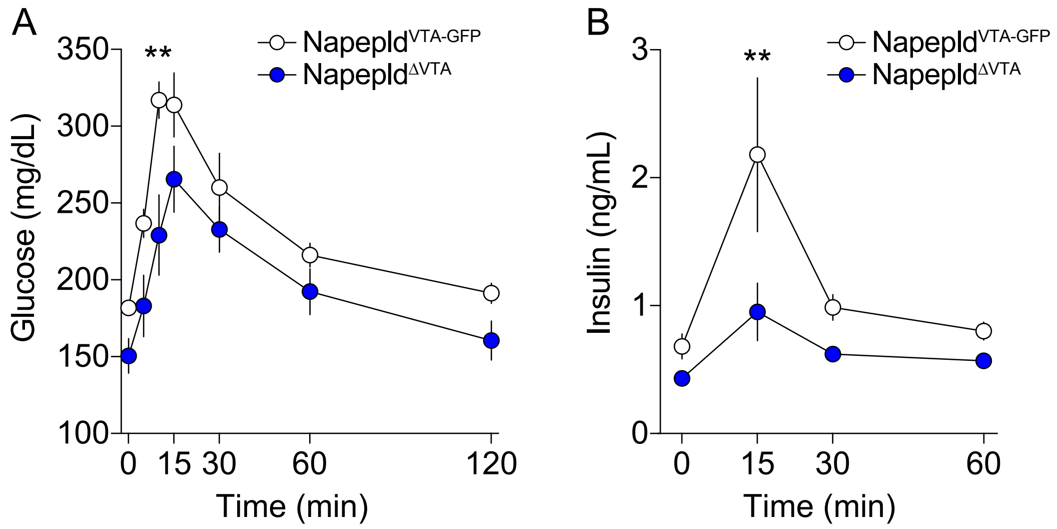
