## Supplementary material for "NAPE-PLD in the ventral tegmental area regulates reward events, feeding and energy homeostasis": Suppl. Table 1

### Supplementary Table 1

| Statistics of Figure 1 | | | | | | |
|---|---|---|---|---|---|---|
| **Figure panels** | | **n** | **Statistical analysis** | | **F-value** | **p-value** |
| Fig 1A | Number of Rewards (FR1) | Napepld$^{f/f}$ n=11, Napepld$^{\Delta CNS}$ n=12 | 2-way ANOVA | interaction | $F_{(3, 63)} = 4.53$ | p = 0.0061 |
| | | | | time | $F_{(3, 63)} = 25.74$ | p < 0.0001 |
| | | | | groups | $F_{(1, 21)} = 5.444$ | p = 0.0297 |
| Fig 1A$^{1}$ | Number of Active Lever Presses (FR1) | Napepld$^{f/f}$ n=11, Napepld$^{\Delta CNS}$ n=12 | 2-way ANOVA | interaction | $F_{(3, 63)} = 2.933$ | p = 0.0402 |
| | | | | time | $F_{(3, 63)} = 16.22$ | p < 0.0001 |
| | | | | groups | $F_{(1, 21)} = 5.842$ | p = 0.0248 |
| Fig 1A$^{2}$ | % of Discrimination (Act/Inact) (FR1) | Napepld$^{f/f}$ n=11, Napepld$^{\Delta CNS}$ n=12 | 2-way ANOVA | interaction | $F_{(3, 63)} = 0.6749$ | p = 0.5707 |
| | | | | time | $F_{(3, 63)} = 1.602$ | p = 0.1978 |
| | | | | groups | $F_{(1, 21)} = 0.0011$ | p = 0.9730 |
| Fig 1B | Number of Rewards (PR) | Napepld$^{f/f}$ n=11, Napepld$^{\Delta CNS}$ n=12 | 2-way ANOVA | interaction | $F_{(3, 63)} = 3.025$ | p = 0.0360 |
| | | | | time | $F_{(3, 63)} = 20.5$ | p < 0.0001 |
| | | | | groups | $F_{(1, 21)} = 7.444$ | p = 0.0126 |
| Fig 1B$^{1}$ | Number of Active Lever Presses (PR) | Napepld$^{f/f}$ n=11, Napepld$^{\Delta CNS}$ n=12 | 2-way ANOVA | interaction | $F_{(3, 63)} = 4.854$ | p = 0.0042 |
| | | | | time | $F_{(3, 63)} = 13.42$ | p < 0.0001 |
| | | | | groups | $F_{(1, 21)} = 7.284$ | p = 0.0134 |
| Fig 1B$^{2}$ | % of Discrimination (Act/Inact) (PR) | Napepld$^{f/f}$ n=11, Napepld$^{\Delta CNS}$ n=12 | 2-way ANOVA | interaction | $F_{(3, 63)} = 1.761$ | p = 0.1637 |
| | | | | time | $F_{(3, 63)} = 8.06$ | p < 0.0001 |
| | | | | groups | $F_{(1, 21)} = 0.0085$ | p = 0.9272 |
| Fig 1C | AEA | Napepld$^{f/f}$ n=4, Napepld$^{\Delta CNS}$ n=5 | 2-way ANOVA | interaction | $F_{(3, 28)} = 5.771$ | p = 0.00330 |
| | | | | time | $F_{(3, 28)} = 69.36$ | p < 0.0001 |
| | | | | groups | $F_{(1, 28)} = 15.22$ | p = 0.0005 |
| Fig 1D | OEA | | 2-way ANOVA | interaction | $F_{(3, 28)} = 3.766$ | p = 0.0218 |
| | | | | structures | $F_{(3, 28)} = 157.80$ | p < 0.0001 |
| | | | | groups | $F_{(1, 28)} = 13.01$ | p = 0.0012 |
| Fig 1E | PEA | | 2-way ANOVA | interaction | $F_{(3, 28)} = 6.444$ | p = 0.0019 |

| | | | | | | |
|---|---|---|---|---|---|---|
| | | | | structures | $F_{(3, 28)}$ = 125.9 | p < 0.0001 |
| | | | | groups | $F_{(1, 28)}$ = 12.97 | p = 0.0012 |
| Fig 1F | SEA | | 2-way ANOVA | interaction | $F_{(3, 28)}$ = 6.195 | p = 0.0023 |
| | | | | structures | $F_{(3, 28)}$ = 57.94 | p < 0.0001 |
| | | | | groups | $F_{(1, 28)}$ = 25.81 | p < 0.0001 |
| Fig 1G | LEA | | 2-way ANOVA | interaction | $F_{(3, 28)}$ = 7.621 | p = 0.0007 |
| | | | | structures | $F_{(3, 28)}$ = 56.8 | p < 0.0001 |
| | | | | groups | $F_{(1, 28)}$ = 38.34 | p < 0.0001 |
| Fig 1H | DEA | | 2-way ANOVA | interaction | $F_{(3, 28)}$ = 5.889 | p = 0.0030 |
| | | | | structures | $F_{(3, 28)}$ = 60.57 | p < 0.0001 |
| | | | | groups | $F_{(1, 8)}$ = 21.52 | p < 0.0001 |
| Fig 1I | 2-AG | | 2-way ANOVA | interaction | $F_{(3, 28)}$ = 1.296 | p = 0.2953 |
| | | | | structures | $F_{(3, 28)}$ = 57.9 | p < 0.0001 |
| | | | | groups | $F_{(1, 28)}$ = 2.236 | p = 0.1460 |

| Statistics of Figure 3 | | | | | | |
|---|---|---|---|---|---|---|
| **Figure panels** | | **n** | **Statistical analysis** | | **F-value** | **p-value** |
| Fig 3B | Number of Rewards (FR1) | $Napepld^{VTA\text{-}GFP}$ n=18, $Napepld^{\Delta VTA}$ n=16 | 2-way ANOVA | interaction | $F_{(2, 64)}$ = 0.5585 | p = 0.5748 |
| | | | | time | $F_{(2, 64)}$ = 15. 86 | p < 0.0001 |
| | | | | groups | $F_{(1, 32)}$ = 2.704 | p = 0.0003 |
| | Active Lever Presses (FR1) | $Napepld^{VTA\text{-}GFP}$ n=18, $Napepld^{\Delta VTA}$ n=16 | 2-way ANOVA | interaction | $F_{(2, 64)}$ = 0.9086 | p = 0.4082 |
| | | | | time | $F_{(2, 64)}$ = 7.712 | p = 0.0010 |
| | | | | groups | $F_{(1, 32)}$ = 7.939 | p = 0.0082 |
| | % of Discrimination (Act/Inact) (FR1) | $Napepld^{VTA\text{-}GFP}$ n=18, $Napepld^{\Delta VTA}$ n=16 | 2-way ANOVA | interaction | $F_{(2, 64)}$ = 0.0054 | p = 0.9946 |
| | | | | time | $F_{(2, 64)}$ = 3.035 | p = 0.0551 |
| | | | | groups | $F_{(1, 32)}$ = 0.0833 | p = 0.7747 |

| | | | | | | |
|---|---|---|---|---|---|---|
| Fig 3C | Number of Rewards (PR) | Napepld$^{VTA-GFP}$ n=18, Napepld$^{\Delta VTA}$ n=16 | 2-way ANOVA | interaction | $F_{(2, 64)} = 0.5873$ | $p = 0.5588$ |
| | | | | time | $F_{(2, 64)} = 7.148$ | $p = 0.0016$ |
| | | | | groups | $F_{(1, 32)} = 9.762$ | $p = 0.0038$ |
| | Active Lever Presses (PR) | Napepld$^{VTA-GFP}$ n=18, Napepld$^{\Delta VTA}$ n=16 | 2-way ANOVA | interaction | $F_{(2, 64)} = 2.719$ | $p = 0.0736$ |
| | | | | time | $F_{(2, 64)} = 6.510$ | $p = 0.0027$ |
| | | | | groups | $F_{(1, 32)} = 8.132$ | $p = 0.0076$ |
| | % of Discrimination (Act/Inact) (PR) | Napepld$^{VTA-GFP}$ n=18, Napepld$^{\Delta VTA}$ n=16 | 2-way ANOVA | interaction | $F_{(2, 64)} = 1.967$ | $p = 0.1483$ |
| | | | | time | $F_{(2, 64)} = 1.378$ | $p = 0.2595$ |
| | | | | groups | $F_{(1, 32)} = 0.6687$ | $p = 0.4196$ |
| Fig 3F | CPP (food restriction) | Napepld$^{VTA-GFP}$ n=8, Napepld$^{\Delta VTA}$ n=8 | 2-way ANOVA | interaction | $F_{(1, 14)} = 0.1543$ | $p = 0.7004$ |
| | | | | time | $F_{(1, 14)} = 0.2688$ | $p = 0.6122$ |
| | | | | groups | $F_{(1, 14)} = 21.81$ | $p = 0.0004$ |
| Fig 3G | CPP (food restriction) | Napepld$^{VTA-GFP}$ n=8, Napepld$^{\Delta VTA}$ n=8 | 2-way ANOVA | interaction | $F_{(1, 14)} = 9.408$ | $p = 0.0084$ |
| | | | | time | $F_{(1, 14)} = 2.192$ | $p = 0.1609$ |
| | | | | groups | $F_{(1, 14)} = 14.78$ | $p = 0.0018$ |
| Fig 3H | T-Maze (learning) | Napepld$^{VTA-GFP}$ n=10, Napepld$^{\Delta VTA}$ n=10 | 2-way ANOVA | interaction | $F_{(4, 56)} = 1.987$ | $p = 0.1090$ |
| | | | | time | $F_{(4, 56)} = 66.64$ | $p < 0.0001$ |
| | | | | groups | $F_{(1, 14)} = 23.44$ | $p = 0.0003$ |
| Fig 3H | T-Maze (reversal) | Napepld$^{VTA-GFP}$ n=10, Napepld$^{\Delta VTA}$ n=10 | 2-way ANOVA | interaction | $F_{(4, 56)} = 3.556$ | $p = 0.0118$ |
| | | | | time | $F_{(4, 56)} = 101.8$ | $p < 0.0001$ |
| | | | | groups | $F_{(1, 14)} = 17.39$ | $p = 0.0009$ |
| Fig 3I | Sucralose preference | Napepld$^{VTA-GFP}$ n=8, Napepld$^{\Delta VTA}$ n=7 | 2-way ANOVA | interaction | $F_{(1, 13)} = 20.25$ | $p = 0.0005$ |
| | | | | time | $F_{(1, 13)} = 110.3$ | $p < 0.0001$ |
| | | | | groups | $F_{(1, 13)} = 14.29$ | $p = 0.0020$ |
| Fig 3J | Sucrose preference | Napepld$^{VTA-GFP}$ n=8, Napepld$^{\Delta VTA}$ n=7 | 2-way ANOVA | interaction | $F_{(1, 13)} = 57.03$ | $p < 0.0001$ |
| | | | | time | $F_{(1, 13)} = 467.9$ | $p < 0.0001$ |
| | | | | groups | $F_{(1, 13)} = 45.54$ | $p < 0.0001$ |
| Fig 3K | Lipids preference | Napepld$^{VTA-GFP}$ n=8, | 2-way ANOVA | interaction | $F_{(1, 13)} = 62,19$ | $p < 0.0001$ |
| | | | | time | $F_{(1, 13)} = 422.4$ | $p < 0.0001$ |

| | | | | | | |
|---|---|---|---|---|---|---|
| | | Napepld$^{\Delta VTA}$ n=7 | | groups | $F_{(1, 13)} = 40.68$ | $p < 0.0001$ |
| Fig 3L | cFos in the NAc | Napepld$^{VTA\text{-}GFP}$ n=6, Napepld$^{\Delta VTA}$ n=6 | Unpaired t-test | | | p = 0.0006 t=4.981, df=10 |

**Statistics of Figure 4**

| Figure panels | | n | Statistical analysis | | F-value | p-value |
|---|---|---|---|---|---|---|
| Fig 4D | Dopamine release (HFD in fasted mice) | Napepld$^{VTA\text{-}GFP}$ n=6, Napepld$^{\Delta VTA}$ n=7 | 2-way ANOVA | interaction | $F_{(2, 22)} = 5.926$ | $p = 0.0087$ |
| | | | | time | $F_{(2, 22)} = 44.92$ | $p < 0.0001$ |
| | | | | groups | $F_{(1, 11)} = 5.274$ | $p = 0.0226$ |
| Fig 4F | Dopamine release (HFD in fed mice) | Napepld$^{VTA\text{-}GFP}$ n=6, Napepld$^{\Delta VTA}$ n=7 | 2-way ANOVA | interaction | $F_{(2, 22)} = 5.308$ | $p = 0.0131$ |
| | | | | time | $F_{(2, 22)} = 47.14$ | $p < 0.0001$ |
| | | | | groups | $F_{(1, 11)} = 8.896$ | $p = 0.0125$ |
| Fig 4H | Dopamine release (cocaine) | Napepld$^{VTA\text{-}GFP}$ n=6, Napepld$^{\Delta VTA}$ n=7 | 2-way ANOVA | interaction | $F_{(3, 33)} = 3.821$ | $p = 0.0187$ |
| | | | | time | $F_{(3, 33)} = 27.65$ | $p < 0.0001$ |
| | | | | groups | $F_{(1, 11)} = 6.822$ | $p = 0.0242$ |
| Fig 4J | Dopamine release (tail suspension) | Napepld$^{VTA\text{-}GFP}$ n=6, Napepld$^{\Delta VTA}$ n=7 | 2-way ANOVA | interaction | $F_{(2, 22)} = 7.161$ | $p = 0.0040$ |
| | | | | time | $F_{(2, 22)} = 29.58$ | $p < 0.0001$ |
| | | | | groups | $F_{(1, 11)} = 7.655$ | $p = 0.0183$ |
| | | | | groups | $F_{(1, 10)} = 0.134$ | $p = 0.7219$ |

**Statistics of Figure 5**

| Figure panels | | n | Statistical analysis | | F-value | p-value |
|---|---|---|---|---|---|---|
| Fig 5A | Body weight (g) | Napepld$^{VTA\text{-}GFP}$ n=14, Napepld$^{\Delta VTA}$ n=18 | Unpaired t-test | | | p = 0.1550 t=1.459, df=30 |
| | Lean body mass (g) | | | | | p = 0.5534 t=0.5994, df=30 |
| | Fat body mass (g) | | | | | p = 0.0729 t=1.859, df=30 |
| Fig 5B | Locomotor activity | Napepld$^{VTA\text{-}GFP}$ n=5, Napepld$^{\Delta VTA}$ n=5 | 2-way ANOVA | interaction | $F_{(96, 768)} = 1,229$ | $p = 0.0773$ |
| | | | | time | $F_{(96, 768)} = 1.605$ | $p = 0.0004$ |

| | | | | | | |
|---|---|---|---|---|---|---|
| | | | | groups | $F_{(1, 8)} = 22.89$ | $p = 0.0014$ |
| Fig 5B[1] | Locomotor activity | $Napepld^{VTA\text{-}GFP}$ n=5, $Napepld^{\Delta VTA}$ n=5 | 2-way ANOVA | interaction | $F_{(1, 8)} = 0.5771$ | $p = 0.4693$ |
| | | | | time | $F_{(1, 8)} = 17.6$ | $p = 0.0030$ |
| | | | | groups | $F_{(1, 8)} = 23.26$ | $p = 0.0013$ |
| Fig 5C | Food intake | $Napepld^{VTA\text{-}GFP}$ n=5, $Napepld^{\Delta VTA}$ n=5 | 2-way ANOVA | interaction | $F_{(96, 768)} = 9{,}378$ | $p < 0.0001$ |
| | | | | time | $F_{(96, 768)} = 320.1$ | $p < 0.0001$ |
| | | | | groups | $F_{(1, 8)} = 23.28$ | $p = 0.0013$ |
| Fig 5C[1] | Food intake | $Napepld^{VTA\text{-}GFP}$ n=5, $Napepld^{\Delta VTA}$ n=5 | 2-way ANOVA | interaction | $F_{(1, 8)} = 32.57$ | $p = 0.0005$ |
| | | | | time | $F_{(1, 8)} = 828.3$ | $p < 0.0001$ |
| | | | | groups | $F_{(1, 8)} = 20.3$ | $p = 0.0020$ |
| Fig 5D | Energy expenditure (EE) | $Napepld^{VTA\text{-}GFP}$ n=5, $Napepld^{\Delta VTA}$ n=5 | 2-way ANOVA | interaction | $F_{(96, 768)} = 1.43$ | $p = 0.0065$ |
| | | | | time | $F_{(96, 768)} = 2.896$ | $p < 0.0001$ |
| | | | | groups | $F_{(1, 8)} = 13.67$ | $p = 0.0061$ |
| Fig 5D[1] | Energy expenditure (EE) | $Napepld^{VTA\text{-}GFP}$ n=5, $Napepld^{\Delta VTA}$ n=5 | 2-way ANOVA | interaction | $F_{(1, 8)} = 1.244$ | $p = 0.2970$ |
| | | | | time | $F_{(1, 8)} = 52.6$ | $p < 0.0001$ |
| | | | | groups | $F_{(1, 8)} = 13.67$ | $p = 0.0061$ |
| Fig 5E | Respiratory Exchange Ratio (RER) | $Napepld^{VTA\text{-}GFP}$ n=5, $Napepld^{\Delta VTA}$ n=5 | 2-way ANOVA | interaction | $F_{(96, 768)} = 2{,}969$ | $p < 0.0001$ |
| | | | | time | $F_{(96, 768)} = 8.308$ | $p < 0.0001$ |
| | | | | groups | $F_{(1, 8)} = 2.02$ | $p = 0.1931$ |
| Fig 5E[1] | Respiratory Exchange Ratio (RER) | $Napepld^{VTA\text{-}GFP}$ n=5, $Napepld^{\Delta VTA}$ n=5 | 2-way ANOVA | interaction | $F_{(1, 8)} = 15.58$ | $p = 0.0043$ |
| | | | | time | $F_{(1, 8)} = 67.74$ | $p < 0.0001$ |
| | | | | groups | $F_{(1, 8)} = 5.308$ | $p = 0.050$ |
| Fig 5F | Fatty acid oxidation (FAO) | $Napepld^{VTA\text{-}GFP}$ n=5, $Napepld^{\Delta VTA}$ n=5 | 2-way ANOVA | interaction | $F_{(96, 768)} = 2.343$ | $p < 0.0001$ |
| | | | | time | $F_{(96, 768)} = 8.108$ | $p < 0.0001$ |
| | | | | groups | $F_{(1, 8)} = 6.77$ | $p = 0.0315$ |
| Fig 5F[1] | Fatty acid oxidation (FAO) | $Napepld^{VTA\text{-}GFP}$ n=5, $Napepld^{\Delta VTA}$ n=5 | 2-way ANOVA | interaction | $F_{(1, 8)} = 0.9988$ | $p = 0.3469$ |
| | | | | time | $F_{(1, 8)} = 27{,}04$ | $p = 0.0008$ |
| | | | | groups | $F_{(1, 8)} = 6.514$ | $p = 0.0341$ |
| Fig 5G | Locomotor activity (fasting/refeeding) | $Napepld^{VTA\text{-}GFP}$ n=6, | 2-way ANOVA | interaction | $F_{(144, 1296)} = 6.35$ | $p < 0.0001$ |
| | | | | time | $F_{(144, 1296)} = 7.64$ | $p < 0.0001$ |

| | | | | | | |
|---|---|---|---|---|---|---|
| | | Napepld$^{\Delta VTA}$ n=5 | | groups | $F_{(1, 9)} = 11.53$ | p = 0.0079 |
| Fig 5H | Food intake (refeeding) | Napepld$^{VTA\text{-}GFP}$ n=6, Napepld$^{\Delta VTA}$ n=5 | Unpaired t-test | | | p = 0.0576 t=2.186, df=9 |
| Fig 5I | Energy expenditure (fasting/refeeding) | Napepld$^{VTA\text{-}GFP}$ n=6, Napepld$^{\Delta VTA}$ n=5 | 2-way ANOVA | interaction | $F_{(144, 1296)} = 3.436$ | p < 0.0001 |
| | | | | time | $F_{(144, 1296)} = 6.392$ | p < 0.0001 |
| | | | | groups | $F_{(1, 9)} = 64.56$ | p < 0.0001 |
| Fig 5J | RER (fasting/refeeding) | Napepld$^{VTA\text{-}GFP}$ n=6, Napepld$^{\Delta VTA}$ n=5 | 2-way ANOVA | interaction | $F_{(144, 1296)} = 2.125$ | p < 0.0001 |
| | | | | time | $F_{(144, 1296)} = 356.9$ | p < 0.0001 |
| | | | | groups | $F_{(1, 9)} = 2,042$ | p = 0.1868 |
| Fig 5K | FAO (fasting/refeeding) | Napepld$^{VTA\text{-}GFP}$ n=6, Napepld$^{\Delta VTA}$ n=5 | 2-way ANOVA | interaction | $F_{(144, 1296)} = 4.413$ | p < 0.0001 |
| | | | | time | $F_{(144, 1296)} = 145.1$ | p < 0.0001 |
| | | | | groups | $F_{(1, 9)} = 34.53$ | p = 0.0002 |
| Fig 5M | Dopamine release (chow pellet in fasted mice) | Napepld$^{VTA\text{-}GFP}$ n=6, Napepld$^{\Delta VTA}$ n=6 | 2-way ANOVA | interaction | $F_{(2, 22)} = 1.101$ | p = 0.3502 |
| | | | | time | $F_{(2, 22)} = 17.35$ | p < 0.0001 |
| | | | | groups | $F_{(1, 11)} = 0.2204$ | p = 0.6479 |

| **Statistics of Figure 6** | | | | | | |
|---|---|---|---|---|---|---|
| **Figure panels** | | **n** | **Statistical analysis** | | **F-value** | **p-value** |
| Fig 6A | Running wheel | Napepld$^{VTA\text{-}GFP}$ n=7, Napepld$^{\Delta VTA}$ n=7 | 2-way ANOVA | interaction | $F_{(4, 48)} = 4.999$ | p = 0.0019 |
| | | | | time | $F_{(4, 48)} = 125.3$ | p < 0.0001 |
| | | | | groups | $F_{(1, 12)} = 5.98$ | p = 0.0309 |
| Fig 6A$^{1}$ | Running wheel | Napepld$^{VTA\text{-}GFP}$ n=7, Napepld$^{\Delta VTA}$ n=7 | 2-way ANOVA | interaction | $F_{(1, 12)} = 9.783$ | p = 0.0087 |
| | | | | time | $F_{(1, 12)} = 280.0$ | p < 0.0001 |
| | | | | groups | $F_{(1, 12)} = 3.042$ | p = 0.1066 |
| Fig 6B | Spontaneous activity | Napepld$^{VTA\text{-}GFP}$ n=10, Napepld$^{\Delta VTA}$ n=7 | 2-way ANOVA | interaction | $F_{(96, 1440)} = 6.913$ | p < 0.0001 |
| | | | | time | $F_{(96, 1440)} = 11.14$ | p < 0.0001 |
| | | | | groups | $F_{(1, 15)} = 59.9$ | p < 0.0001 |
| Fig 6B$^{1}$ | Spontaneous activity | Napepld$^{VTA\text{-}GFP}$ n=10, Napepld$^{\Delta VTA}$ n=7 | 2-way ANOVA | interaction | $F_{(1, 15)} = 14.96$ | p = 0.0015 |
| | | | | time | $F_{(1, 15)} = 24.78$ | p = 0.0002 |
| | | | | groups | $F_{(1, 15)} = 60.4$ | p < 0.0001 |
| Fig 6C | Wheel activity | Napepld$^{VTA\text{-}GFP}$ n=10, | 2-way ANOVA | interaction | $F_{(96, 1440)} = 3.344$ | p < 0.0001 |

| | | | | | | |
|---|---|---|---|---|---|---|
| | | $Napepld^{\Delta VTA}$ n=7 | | time | $F_{(96, 1440)} = 21.18$ | $p < 0.0001$ |
| | | | | groups | $F_{(1, 15)} = 6.033$ | $p = 0.0267$ |
| Fig 6C[1] | Wheel activity | $Napepld^{VTA\text{-}GFP}$ n=10, $Napepld^{\Delta VTA}$ n=7 | 2-way ANOVA | interaction | $F_{(1, 15)} = 16.92$ | $p = 0.0009$ |
| | | | | time | $F_{(1, 15)} = 124.2$ | $p < 0.0001$ |
| | | | | groups | $F_{(1, 15)} = 9.76$ | $p = 0.0070$ |
| Fig 6D | All activities | $Napepld^{VTA\text{-}GFP}$ n=10, $Napepld^{\Delta VTA}$ n=7 | 2-way ANOVA | interaction | $F_{(1, 15)} = 5.847$ | $p = 0.0288$ |
| | | | | time | $F_{(1, 15)} = 125.1$ | $p < 0.0001$ |
| | | | | groups | $F_{(1, 15)} = 2.12$ | $p = 0.1660$ |
| Fig 6E | Energy expenditure | $Napepld^{VTA\text{-}GFP}$ n=10, $Napepld^{\Delta VTA}$ n=7 | 2-way ANOVA | interaction | $F_{(96, 1440)} = 1.282$ | $p = 0.0385$ |
| | | | | time | $F_{(96, 1440)} = 36.71$ | $p < 0.0001$ |
| | | | | groups | $F_{(1, 15)} = 10.15$ | $p = 0.0061$ |
| Fig 6E[1] | Energy expenditure | $Napepld^{VTA\text{-}GFP}$ n=10, $Napepld^{\Delta VTA}$ n=7 | 2-way ANOVA | interaction | $F_{(1, 15)} = 0.1591$ | $p = 0.6956$ |
| | | | | time | $F_{(1, 15)} = 174.9$ | $p < 0.0001$ |
| | | | | groups | $F_{(1, 15)} = 10.21$ | $p = 0.0060$ |
| Fig 6F | Food intake | $Napepld^{VTA\text{-}GFP}$ n=10, $Napepld^{\Delta VTA}$ n=7 | 2-way ANOVA | interaction | $F_{(96, 1440)} = 8.714$ | $p < 0.0001$ |
| | | | | time | $F_{(96, 1440)} = 424.6$ | $p < 0.0001$ |
| | | | | groups | $F_{(1, 15)} = 9.391$ | $p = 0.0079$ |
| Fig 6G | RER | $Napepld^{VTA\text{-}GFP}$ n=10, $Napepld^{\Delta VTA}$ n=7 | 2-way ANOVA | interaction | $F_{(1, 15)} = 3.885$ | $p = 0.0674$ |
| | | | | time | $F_{(1, 15)} = 43.38$ | $p < 0.0001$ |
| | | | | groups | $F_{(1, 15)} = 6.297$ | $p = 0.0005$ |
| Fig 6H | FAO | $Napepld^{VTA\text{-}GFP}$ n=10, $Napepld^{\Delta VTA}$ n=7 | 2-way ANOVA | interaction | $F_{(1, 15)} = 15.18$ | $p = 0.0014$ |
| | | | | time | $F_{(1, 15)} = 2.613$ | $p = 0.1268$ |
| | | | | groups | $F_{(1, 15)} = 7.031$ | $p = 0.0003$ |

| **Statistics of Figure 7** | | | | | | |
|---|---|---|---|---|---|---|
| **Figure panels** | | **n** | **Statistical analysis** | | **F-value** | **p-value** |
| Fig 7A | Body weight (obesity) | $Napepld^{VTA\text{-}GFP}$ n=10, $Napepld^{\Delta VTA}$ n=8 | Unpaired t-test | | | $p = 0.1487$ t=1.517, df=16 |
| | Lean body mass (obesity) | | | | | $p = 0.8180$ t=0.2339, df=16 |
| | Fat body mass (obesity) | | | | | $p = 0.0164$ t=2.68, df=16 |
| Fig 7B | Locomotor activity | $Napepld^{VTA\text{-}GFP}$ n=8, $Napepld^{\Delta VTA}$ n=5 | 2-way ANOVA | interaction | $F_{(1, 11)} = 16.22$ | $p = 0.0020$ |
| | | | | time | $F_{(1, 11)} = 32.49$ | $p = 0.0001$ |
| | | | | groups | $F_{(1, 11)} = 42.94$ | $p < 0.0001$ |
| | Food intake | | | interaction | $F_{(96, 1056)} = 11.38$ | $p < 0.0001$ |

| | | | | | | |
|---|---|---|---|---|---|---|
| Fig 7C | | $Napepld^{VTA-GFP}$ n=8, $Napepld^{\Delta VTA}$ n=5 | 2-way ANOVA | treatment | $F_{(96, 1056)} = 361.1$ | $p < 0.0001$ |
| | | | | groups | $F_{(1, 11)} = 9.033$ | $p = 0.0120$ |
| Fig 7C$^1$ | Food intake | $Napepld^{VTA-GFP}$ n=8, $Napepld^{\Delta VTA}$ n=5 | 2-way ANOVA | interaction | $F_{(1, 11)} = 30.8$ | $p = 0.0002$ |
| | | | | time | $F_{(1, 11)} = 442$ | $p < 0.0001$ |
| | | | | groups | $F_{(1, 11)} = 12.13$ | $p = 0.0051$ |
| Fig 7D | Energy expenditure | $Napepld^{VTA-GFP}$ n=8, $Napepld^{\Delta VTA}$ n=5 | 2-way ANOVA | interaction | $F_{(96, 1056)} = 8.621$ | $p < 0.0001$ |
| | | | | time | $F_{(96, 1056)} = 26.2$ | $p < 0.0001$ |
| | | | | groups | $F_{(1, 11)} = 38.21$ | $p < 0.0001$ |
| Fig 7D$^1$ | Energy expenditure | $Napepld^{VTA-GFP}$ n=8, $Napepld^{\Delta VTA}$ n=5 | 2-way ANOVA | interaction | $F_{(1, 11)} = 20.84$ | $p = 0.0008$ |
| | | | | time | $F_{(1, 11)} = 86.5$ | $p < 0.0001$ |
| | | | | groups | $F_{(1, 11)} = 37.91$ | $p < 0.0001$ |
| Fig 7E | FAO | $Napepld^{VTA-GFP}$ n=8, $Napepld^{\Delta VTA}$ n=5 | 2-way ANOVA | interaction | $F_{(96, 1056)} = 2.319$ | $p < 0.0001$ |
| | | | | time | $F_{(96, 1056)} = 8.215$ | $p < 0.0001$ |
| | | | | groups | $F_{(1, 11)} = 11.78$ | $p = 0.0056$ |
| Fig 7E$^1$ | FAO | $Napepld^{VTA-GFP}$ n=8, $Napepld^{\Delta VTA}$ n=5 | 2-way ANOVA | interaction | $F_{(1, 11)} = 1.169$ | $p = 0.3027$ |
| | | | | time | $F_{(1, 11)} = 27.39$ | $p = 0.0003$ |
| | | | | groups | $F_{(1, 11)} = 11.66$ | $p = 0.0058$ |
| Fig 7F | RER | $Napepld^{VTA-GFP}$ n=8, $Napepld^{\Delta VTA}$ n=5 | 2-way ANOVA | interaction | $F_{(1, 11)} = 3.64$ | $p = 0.0829$ |
| | | | | time | $F_{(1, 11)} = 5.968$ | $p = 0.0327$ |
| | | | | groups | $F_{(1, 11)} = 0.2319$ | $p = 0.6395$ |

| **Statistics of Suppl. Figure 1** | | | | | | |
|---|---|---|---|---|---|---|
| **Figure panels** | | **n** | **Statistical analysis** | | **F-value** | **p-value** |
| SFig 1A | Number of Rewards (FR1) | $Napepld^{+/+}$ n=8, $Napepld^{KO}$ n=8 | 2-way ANOVA | interaction | $F_{(3, 42)} = 0,5499$ | $p = 0.6510$ |
| | | | | time | $F_{(3, 42)} = 17,83$ | $p < 0.0001$ |
| | | | | groups | $F_{(1, 14)} = 0,0267$ | $p = 0.8724$ |
| | Active lever presses (FR1) | | 2-way ANOVA | interaction | $F_{(3, 42)} = 0.5366$ | $p = 0.6598$ |
| | | | | time | $F_{(3, 42)} = 11.49$ | $p < 0.0001$ |
| | | | | groups | $F_{(1, 14)} = 0.4121$ | $p = 0.5313$ |
| | % of Discrimination (FR1) | | 2-way ANOVA | interaction | $F_{(3, 42)} = 1.708$ | $p = 0.1800$ |
| | | | | time | $F_{(3, 42)} = 4.775$ | $p = 0.0059$ |
| | | | | groups | $F_{(1, 14)} = 0.03782$ | $p = 0.8486$ |

| | | | | | | |
|---|---|---|---|---|---|---|
| SFig 1B | Number of Rewards (PR) | Napepld$^{+/+}$ n=8, Napepld$^{KO}$ n=8 | 2-way ANOVA | interaction | $F_{(3, 42)} = 0.3998$ | $p = 0.7538$ |
| | | | | time | $F_{(3, 42)} = 29.09$ | $p < 0.0001$ |
| | | | | groups | $F_{(1, 14)} = 0.0941$ | $p = 0.7636$ |
| | Active lever presses (PR) | | 2-way ANOVA | interaction | $F_{(3, 42)} = 0.6919$ | $p = 0.5621$ |
| | | | | time | $F_{(3, 42)} = 17.59$ | $p < 0.0001$ |
| | | | | groups | $F_{(1, 14)} = 0,06497$ | $p = 0.8025$ |
| | % of Discrimination (PR) | | 2-way ANOVA | interaction | $F_{(3, 42)} = 2.669$ | $p =0.0598$ |
| | | | | time | $F_{(3, 42)} = 7.503$ | $p = 0.0004$ |
| | | | | groups | $F_{(1, 14)} = 0.1729$ | $p = 0.6839$ |
| SFig 1C | Number of Rewards (FR1) | Nestin$^{Cre-/-}$ n=5, Nestin$^{Cre+/-}$ n=6 | 2-way ANOVA | interaction | $F_{(3, 27)} = 0.8579$ | $p = 0.4748$ |
| | | | | time | $F_{(3, 27)} = 23.13$ | $p < 0.0001$ |
| | | | | groups | $F_{(1, 9)} = 0.004349$ | $p = 0.9489$ |
| | Active lever presses (FR1) | | 2-way ANOVA | interaction | $F_{(3, 27)} = 0.3844$ | $p = 0.7650$ |
| | | | | time | $F_{(3, 27)} = 9.225$ | $p = 0.0002$ |
| | | | | groups | $F_{(1, 9)} = 0.09229$ | $p = 0.7682$ |
| | % of Discrimination (FR1) | | 2-way ANOVA | interaction | $F_{(3, 27)} = 0.987$ | $p = 0.4136$ |
| | | | | time | $F_{(3, 27)} = 0.9717$ | $p = 0.4205$ |
| | | | | groups | $F_{(1, 9)} = 1.484$ | $p = 0.2541$ |
| SFig 1D | Number of Rewards (PR) | Nestin$^{Cre-/-}$ n=5, Nestin$^{Cre+/-}$ n=6 | 2-way ANOVA | interaction | $F_{(3, 27)} = 0.8336$ | $p = 0.4872$ |
| | | | | time | $F_{(3, 27)} = 6.237$ | $p = 0.0023$ |
| | | | | groups | $F_{(1, 9)} = 0.02488$ | $p = 0.8782$ |
| | Active lever presses (PR) | | 2-way ANOVA | interaction | $F_{(3, 27)} = 0.5632$ | $p = 0.6440$ |
| | | | | time | $F_{(3, 27)} = 5.384$ | $p = 0.0049$ |
| | | | | groups | $F_{(1, 9)} = 0.3782$ | $p = 0.5538$ |
| | % of Discrimination (PR) | | 2-way ANOVA | interaction | $F_{(3, 27)} = 4.498$ | $p = 0.0110$ |
| | | | | time | $F_{(3, 27)} = 1.25$ | $p = 0.3111$ |
| | | | | groups | $F_{(1, 9)} = 0.05107$ | $p = 0.8263$ |
| SFig 1E | Number of Rewards (FR1) | Napepld$^{f/f}$ n=6, Napepld$^{\Delta IEC}$ n=6 | 2-way ANOVA | interaction | $F_{(3, 30)} = 0.1818$ | $p = 0.9079$ |
| | | | | time | $F_{(3, 30)} = 16.81$ | $p < 0.0001$ |

| | | | | | | |
|---|---|---|---|---|---|---|
| | | | | groups | $F_{(1, 10)} = 0.3784$ | p = 0.5522 |
| | Active lever presses (FR1) | | 2-way ANOVA | interaction | $F_{(3, 30)} = 0.04352$ | p = 0.9877 |
| | | | | time | $F_{(3, 30)} = 12.16$ | p < 0.0001 |
| | | | | groups | $F_{(1, 10)} = 1.436$ | p = 0.2583 |
| | % of Discrimination (FR1) | | 2-way ANOVA | interaction | $F_{(3, 30)} = 0.1166$ | p = 0.9497 |
| | | | | time | $F_{(3, 30)} = 0.6663$ | p = 0.5793 |
| | | | | groups | $F_{(1, 10)} = 0.2446$ | p = 0.6316 |
| SFig 1F | Number of Rewards (PR) | Napepld$^{f/f}$ n=6, Napepld$^{\Delta IEC}$ n=6 | 2-way ANOVA | interaction | $F_{(3, 30)} = 0.5034$ | p = 0.6828 |
| | | | | time | $F_{(3, 30)} = 37.88$ | p < 0.0001 |
| | | | | groups | $F_{(1, 10)} = 0.3644$ | p = 0.5595 |
| | Active lever presses (PR) | | 2-way ANOVA | interaction | $F_{(3, 30)} = 0.5376$ | p = 0.6601 |
| | | | | time | $F_{(3, 30)} = 20.34$ | p < 0.0001 |
| | | | | groups | $F_{(1, 10)} = 0.235$ | p = 0.6383 |
| | % of Discrimination (PR) | | 2-way ANOVA | interaction | $F_{(3, 30)} = 0.3135$ | p = 0.8154 |
| | | | | time | $F_{(3, 30)} = 16.44$ | p < 0.0001 |
| | | | | groups | $F_{(1, 10)} = 0.03408$ | p = 0.8572 |

**Statistics of Suppl. Figure 2**

| Figure panels | | n | Statistical analysis | F-value | p-value |
|---|---|---|---|---|---|
| SFig 2A | Linoleic acid | Napepld$^{f/f}$ n=4, Napepld$^{\Delta CNS}$ n=5 | Unpaired t-test | | p = 0.6256, t=0.5102, df=7 |
| SFig 2B | Arachidonic acid | | | | p = 0.5357, t=0.6513, df=7 |
| SFig 2C | Oleic acid | | | | p = 0.4907, t=0.7271, df=7 |

**Statistics of Suppl. Figure 3**

| Figure panels | | n | Statistical analysis | | F-value | p-value |
|---|---|---|---|---|---|---|
| SFig 3A | Active lever presses (FR1) | Napepld$^{VTA-GFP}$ n=9, Napepld$^{\Delta VTA}$ n=9 | 2-way ANOVA | interaction | $F_{(4, 64)} = 1.39$ | p = 0.2475 |
| | | | | time | $F_{(4, 64)} = 3.633$ | p = 0.0099 |
| | | | | groups | $F_{(1, 16)} = 19.18$ | p = 0.0005 |
| | Number of Rewards (FR1) | | 2-way ANOVA | interaction | $F_{(4, 64)} = 1.095$ | p = 0.3667 |
| | | | | time | $F_{(4, 64)} = 6.529$ | p = 0.0002 |
| | | | | groups | $F_{(1, 16)} = 30.57$ | p < 0.0001 |
| SFig 3B | Active lever presses (FR5) | Napepld$^{VTA-GFP}$ n=9, Napepld$^{\Delta VTA}$ n=9 | 2-way ANOVA | interaction | $F_{(4, 64)} = 0.272$ | p = 0.8950 |
| | | | | time | $F_{(4, 64)} = 2.33$ | p = 0.0654 |
| | | | | groups | $F_{(1, 16)} = 29.65$ | p < 0.0001 |
| | Number of Rewards (FR5) | | | interaction | $F_{(4, 64)} = 0.4597$ | p = 0.7650 |

| | | | | | | |
|---|---|---|---|---|---|---|
| | | | 2-way ANOVA | time | $F_{(4, 64)} = 2.748$ | $p = 0.0357$ |
| | | | | groups | $F_{(1, 16)} = 31.44$ | $p < 0.0001$ |
| SFig 3C | Active lever presses (PR) | Napepld$^{VTA\text{-}GFP}$ n=9, Napepld$^{\Delta VTA}$ n=9 | 2-way ANOVA | interaction | $F_{(2, 32)} = 2.322$ | $p = 0.1144$ |
| | | | | time | $F_{(2, 32)} = 3.69$ | $p = 0.0361$ |
| | | | | groups | $F_{(1, 16)} = 18.67$ | $p = 0.0005$ |
| | Number of Rewards (PR) | | 2-way ANOVA | interaction | $F_{(2, 32)} = 0.9863$ | $p = 0.3840$ |
| | | | | time | $F_{(2, 32)} = 3.65$ | $p = 0.0373$ |
| | | | | groups | $F_{(1, 16)} = 20.67$ | $p = 0.0003$ |
| SFig 3D | Active lever presses (PR, fed) | Napepld$^{VTA\text{-}GFP}$ n=9, Napepld$^{\Delta VTA}$ n=9 | 2-way ANOVA | interaction | $F_{(12, 192)} = 9.845$ | $p < 0.0001$ |
| | | | | time | $F_{(12, 192)} = 85.62$ | $p < 0.0001$ |
| | | | | groups | $F_{(1, 16)} = 10.92$ | $p = 0.0045$ |
| | Number of Rewards (PR, fed) | | 2-way ANOVA | interaction | $F_{(12, 192)} = 7,476$ | $p < 0.0001$ |
| | | | | time | $F_{(12, 192)} = 307,9$ | $p < 0.0001$ |
| | | | | groups | $F_{(1, 16)} = 12,75$ | $p = 0.0025$ |
| SFig 3E | Active lever presses (FR1) | Napepld$^{VTA\text{-}GFP}$ n=8, Napepld$^{\Delta VTA}$ n=7 | 2-way ANOVA | interaction | $F_{(4, 52)} = 2.765$ | $p = 0.0369$ |
| | | | | time | $F_{(4, 52)} = 13.62$ | $p < 0.0001$ |
| | | | | groups | $F_{(1, 13)} = 3.212$ | $p = 0.0964$ |
| | Number of Rewards (FR1) | | 2-way ANOVA | interaction | $F_{(4, 52)} = 1,625$ | $p = 0.1819$ |
| | | | | time | $F_{(4, 52)} = 11.96$ | $p < 0.0001$ |
| | | | | groups | $F_{(1, 13)} = 4.306$ | $p = 0.05$ |
| SFig 3F | Active lever presses (FR5) | Napepld$^{VTA\text{-}GFP}$ n=8, Napepld$^{\Delta VTA}$ n=7 | 2-way ANOVA | interaction | $F_{(4, 52)} = 1.043$ | $p = 0.3942$ |
| | | | | time | $F_{(4, 52)} = 4.323$ | $p = 0.0043$ |
| | | | | groups | $F_{(1, 13)} = 5.867$ | $p = 0.0308$ |
| | Number of Rewards (FR5) | | 2-way ANOVA | interaction | $F_{(4, 52)} = 0.8845$ | $p = 0.4797$ |
| | | | | time | $F_{(4, 52)} = 4.105$ | $p = 0.0058$ |
| | | | | groups | $F_{(1, 13)} = 5.294$ | $p = 0.0386$ |
| SFig 3G | Active lever presses (PR) | Napepld$^{VTA\text{-}GFP}$ n=8, Napepld$^{\Delta VTA}$ n=7 | 2-way ANOVA | interaction | $F_{(2, 26)} = 0.3456$ | $p = 0.7110$ |
| | | | | time | $F_{(2, 26)} = 2.512$ | $p = 0.1006$ |
| | | | | groups | $F_{(1, 13)} = 6.703$ | $p = 0.0225$ |
| | Number of Rewards (PR) | | 2-way ANOVA | interaction | $F_{(2, 26)} = 0.3185$ | $p = 0.7300$ |
| | | | | time | $F_{(2, 26)} = 5.29$ | $p = 0.0118$ |
| | | | | groups | $F_{(1, 13)} = 6.494$ | $p = 0.0243$ |
| SFig 3H | Active lever presses (PR, fed) | Napepld$^{VTA\text{-}GFP}$ n=8, | 2-way ANOVA | interaction | $F_{(12, 156)} = 6.521$ | $p < 0.0001$ |

| | | | | | | |
|---|---|---|---|---|---|---|
| | | Napepld$^{\Delta VTA}$ n=7 | | time | $F_{(12, 156)} = 117.5$ | $p < 0.0001$ |
| | | | | groups | $F_{(1, 13)} = 6.102$ | $p = 0.0281$ |
| | Number of Rewards (PR, fed) | | 2-way ANOVA | interaction | $F_{(12, 156)} = 3.448$ | $p = 0.0002$ |
| | | | | time | $F_{(12, 156)} = 365.5$ | $p < 0.0001$ |
| | | | | groups | $F_{(1, 13)} = 4.155$ | $p = 0.0624$ |

**Statistics of Suppl. Figure 4**

| Figure panels | | n | Statistical analysis | | F-value | p-value |
|---|---|---|---|---|---|---|
| SFig 4A | Body weight (males) | Napepld$^{VTA\text{-}GFP}$ n=9, Napepld$^{\Delta VTA}$ n=9 | 2-way ANOVA | interaction | $F_{(1, 16)} = 0.6336$ | $p = 0.4377$ |
| | | | | time | $F_{(1, 16)} = 143.7$ | $p < 0.0001$ |
| | | | | groups | $F_{(1, 16)} = 2.941$ | $p = 0.1056$ |
| SFig 4B | Body weight (females) | Napepld$^{VTA\text{-}GFP}$ n=8, Napepld$^{\Delta VTA}$ n=7 | 2-way ANOVA | interaction | $F_{(1, 13)} = 0.1895$ | $p = 0.6705$ |
| | | | | time | $F_{(1, 13)} = 120.7$ | $p < 0.0001$ |
| | | | | groups | $F_{(1, 13)} = 0.5585$ | $p = 0.4682$ |

**Statistics of Suppl. Figure 5**

| Figure panels | | n | Statistical analysis | F-value | p-value |
|---|---|---|---|---|---|
| SFig 5C | Body weight | Napepld$^{VTA\text{-}GFP}$ n=8, Napepld$^{\Delta VTA}$ n=9 | Unpaired t-test | | $p = 0.0038$ $t=3.416$, $df=15$ |

**Statistics of Suppl. Figure 6**

| Figure panels | | n | Statistical analysis | | F-value | p-value |
|---|---|---|---|---|---|---|
| SFig 6A | Glucose (OGTT) | Napepld$^{VTA\text{-}GFP}$ n=7, Napepld$^{\Delta VTA}$ n=6 | 2-way ANOVA | interaction | $F_{(6, 66)} = 1.624$ | $p = 0.1544$ |
| | | | | time | $F_{(6, 66)} = 28.65$ | $p < 0.0001$ |
| | | | | groups | $F_{(1, 11)} = 8.518$ | $p = 0.0140$ |
| SFig 6B | Insulin (OGTT) | Napepld$^{VTA\text{-}GFP}$ n=7, Napepld$^{\Delta VTA}$ n=6 | 2-way ANOVA | interaction | $F_{(3, 33)} = 2.297$ | $p = 0.0958$ |
| | | | | time | $F_{(3, 33)} = 8.298$ | $p = 0.0003$ |
| | | | | groups | $F_{(1, 11)} = 5.252$ | $p = 0.0426$ |
