## Supplementary material for "NAPE-PLD in the ventral tegmental area regulates reward events, feeding and energy homeostasis": Suppl. Table 2

Results are shown are mean ± SEM.
BDL: Below Detection Limits
Unit: nmol/gram
Statistics: unpaired Student's t-test

| | | | | | | | |
|---|---|---|---|---|---|---|---|
| ***N*-acyl alanine** | | | | ***N*-acyl GABA** | | | |
| | **Napepld$^{f/f}$ (n=4)** | **Napepld$^{\Delta CNS}$ (n=5)** | **p value** | | **Napepld$^{f/f}$ (n=4)** | **Napepld$^{\Delta CNS}$ (n=5)** | **p value** |
| *N*-palmitoyl alanine | $8.7^{e-3} \pm 1.745^{e-3}$ | $8.96^{e-3} \pm 1.668^{e-3}$ | 0.9179 | *N*-palmitoyl GABA | $1.388^{e-2} \pm 1.819^{e-3}$ | $1.4^{e-2} \pm 1.0^{e-3}$ | 0.9509 |
| *N*-stearoyl alanine | $5.235^{e-4} \pm 6.099^{e-5}$ | $4.132^{e-4} \pm 3.86^{e-5}$ | 0.1549 | *N*-stearoyl GABA | $1.347^{e-2} \pm 1.414^{e-3}$ | $1.215^{e-2} \pm 1.403^{e-3}$ | 0.5366 |
| *N*-oleoyl alanine | $2.835^{e-3} \pm 5.4769^{e-4}$ | $2.908^{e-3} \pm 5.425^{e-4}$ | 0.9296 | *N*-oleoyl GABA | $5.61^{e-3} \pm 5.852^{e-4}$ | $5.57^{e-3} \pm 5.417^{e-4}$ | 0.9616 |
| *N*-linoleoyl alanine | BDL | BDL | | *N*-linoleoyl GABA | BDL | BDL | |
| *N*-arachidonoyl alanine | $7.69^{e-4} \pm 2.835^{e-5}$ | $6.614^{e-4} \pm 8.491^{e-5}$ | 0.3151 | *N*-arachidonoyl GABA | $2.425^{e-2} \pm 2.485^{e-3}$ | $2.72^{e-2} \pm 3.521^{e-3}$ | 0.5375 |
| *N*-docosahexaenoyl alanine | BDL | BDL | | *N*-docosahexaenoyl GABA | $1.66^{e-3} \pm 1.358^{e-4}$ | $2.06^{e-3} \pm 2.005^{e-4}$ | 0.1632 |
| ***N*-acyl glycine** | | | | ***N*-acyl leucine** | | | |
| | **Napepld$^{f/f}$ (n=4)** | **Napepld$^{\Delta CNS}$ (n=5)** | **p value** | | **Napepld$^{f/f}$ (n=4)** | **Napepld$^{\Delta CNS}$ (n=5)** | **p value** |
| *N*-palmitoyl glycine | $4.28^{e-2} \pm 1.334^{e-2}$ | $3.2^{e-2} \pm 4.219^{e-3}$ | 0.4215 | *N*-palmitoyl leucine | $2.305^{e-3} \pm 4.122^{e-4}$ | $2.0^{e-3} \pm 3.286^{e-4}$ | 0.5756 |
| *N*-stearoyl glycine | $2.74^{e-2} \pm 2.392^{e-3}$ | $2.626^{e-2} \pm 5.451^{e-3}$ | 0.8663 | *N*-stearoyl leucine | $7.098^{e-4} \pm 1.046^{e-5}$ | $5.754^{e-4} \pm 3.328^{e-5}$ | 0.2182 |
| *N*-oleoyl glycine | $3.675^{e-2} \pm 4.384^{e-3}$ | $3.604^{e-2} \pm 4.116^{e-3}$ | 0.9099 | *N*-oleoyl leucine | $3.4^{e-4} \pm 6.418^{e-5}$ | $3.75^{e-4} \pm 8.246^{e-5}$ | 0.7579 |
| *N*-linoleoyl glycine | BDL | BDL | | *N*-linoleoyl leucine | BDL | BDL | |
| *N*-arachidonoyl glycine | $7.978^{e-2} \pm 1.664^{e-3}$ | $9.47^{e-2} \pm 9.716^{e-3}$ | 0.2212 | *N*- arachidonoyl leucine | BDL | BDL | |
| *N*-docosahexaenoyl glycine | $3.348^{e-2} \pm 3.954^{e-3}$ | $3.274^{e-2} \pm 4.18^{e-3}$ | 0.9040 | *N*-docosahexaenoyl leucine | BDL | BDL | |
| ***N*-acyl methionine** | | | | ***N*-acyl phenylalanine** | | | |
| | **Napepld$^{f/f}$ (n=4)** | **Napepld$^{\Delta CNS}$ (n=5)** | **p value** | | **Napepld$^{f/f}$ (n=4)** | **Napepld$^{\Delta CNS}$ (n=5)** | **p value** |
| *N*-palmitoyl methionine | BDL | BDL | | *N*-palmitoyl phenylalanine | $7.013^{e-3} \pm 1.133^{e-3}$ | $5.86^{e-3} \pm 4.226^{e-4}$ | 0.3315 |
| *N*-stearoyl methionine | BDL | BDL | | *N*-stearoyl phenylalanine | $2.625^{e-3} \pm 5.794^{e-4}$ | $2.622^{e-3} \pm 2.246^{e-4}$ | 0.9959 |
| *N*-oleoyl methionine | BDL | BDL | | *N*-oleoyl phenylalanine | $2.633^{e-3} \pm 4.124^{e-4}$ | $2.35^{e-3} \pm 1.413^{e-4}$ | 0.4988 |
| *N*-linoleoyl methionine | BDL | BDL | | *N*-linoleoyl phenylalanine | BDL | BDL | |
| *N*-arachidonoyl methionine | BDL | BDL | | *N*-arachidonoyl phenylalanine | $3.48^{e-3} \pm 6.536^{e-4}$ | $3.342^{e-3} \pm 4.824^{e-4}$ | 0.8668 |
| *N*-docosahexaenoyl methionine | BDL | BDL | | *N*-docosahexaenoyl phenylalanine | $2.215^{e-3} \pm 2.602^{e-4}$ | $2.442^{e-3} \pm 2.949^{e-4}$ | 0.5927 |
| ***N*-acyl proline** | | | | ***N*-acyl serine** | | | |
| | **Napepld$^{f/f}$ (n=4)** | **Napepld$^{\Delta CNS}$ (n=5)** | **p value** | | **Napepld$^{f/f}$ (n=4)** | **Napepld$^{\Delta CNS}$ (n=5)** | **p value** |
| *N*-palmitoyl proline | BDL | BDL | | *N*-palmitoyl serine | $1.613^{e-1} \pm 1.174^{e-3}$ | $1.44^{e-1} \pm 1.208^{e-2}$ | 0.3479 |
| *N*-stearoyl proline | BDL | BDL | | *N*-stearoyl serine | $6.2^{e-2} \pm 1.252^{e-3}$ | $6.132^{e-2} \pm 1.099^{e-2}$ | 0.9685 |

| | | | | | | | |
|---|---|---|---|---|---|---|---|
| *N*-oleoyl proline | BDL | BDL | | *N*-oleoyl serine | $4.013^{e-1} \pm 8.24^{e-2}$ | $3.656^{e-1} \pm 8.046^{e-2}$ | 0.7684 |
| *N*-linoleoyl proline | BDL | BDL | | *N*-linoleoyl serine | $2.103^{e-1} \pm 4.693^{e-2}$ | $1.922^{e-1} \pm 4.385^{e-2}$ | 0.7880 |
| *N*-arachidonoyl proline | BDL | BDL | | *N*-arachidonoyl serine | $6.598^{e-2} \pm 7.228^{e-3}$ | $4.534^{e-2} \pm 3.661^{e-3}$ | 0.0297 |
| *N*-docosahexaenoyl proline | BDL | BDL | | *N*-docosahexaenoyl serine | $5.6^{e-2} \pm 7.268^{e-3}$ | $4.878^{e-2} \pm 1.258^{e-2}$ | 0.6581 |
| ***N*-acyl tryptophan** | | | | ***N*-acyl tyrosine** | | | |
| | **Napepld$^{f/f}$ (n=4)** | **Napepld$^{\Delta CNS}$ (n=5)** | **p value** | | **Napepld$^{f/f}$ (n=4)** | **Napepld$^{\Delta CNS}$ (n=5)** | **p value** |
| *N*-palmitoyl tryptophan | BDL | BDL | | *N*-palmitoyl tyrosine | $4.908^{e-3} \pm 8.214^{e-4}$ | $3.8^{e-3} \pm 4.427^{e-4}$ | 0.2480 |
| *N*-stearoyl tryptophan | BDL | BDL | | *N*-stearoyl tyrosine | BDL | BDL | |
| *N*-oleoyl tryptophan | BDL | BDL | | *N*-oleoyl tyrosine | $1.024^{e-3} \pm 2.026^{e-4}$ | $1.102^{e-3} \pm 7.399^{e-5}$ | 0.7027 |
| *N*-linoleoyl tryptophan | BDL | BDL | | *N*-linoleoyl tyrosine | BDL | BDL | |
| *N*-arachidonoyl tryptophan | BDL | BDL | | *N*-arachidonoyl tyrosine | $1.284^{e-3} \pm 2.269^{e-4}$ | $1.717^{e-3} \pm 4.783^{e-4}$ | 0.4783 |
| *N*-docosahexaenoyl tryptophan | BDL | BDL | | *N*-docosahexaenoyl tyrosine | $2.008^{e-3} \pm 4.811^{e-4}$ | $1.707^{e-3} \pm 4.851^{e-4}$ | 0.6781 |
| ***N*-acyl valine** | | | | **Prostaglandins** | | | |
| | **Napepld$^{f/f}$ (n=4)** | **Napepld$^{\Delta CNS}$ (n=5)** | **p value** | | **Napepld$^{f/f}$ (n=4)** | **Napepld$^{\Delta CNS}$ (n=5)** | **p value** |
| *N*-palmitoyl valine | $1.508^{e-3} \pm 7.499^{e-5}$ | $1.174^{e-3} \pm 2.03^{e-4}$ | 0.2063 | $PGE_2$ | $1.508 \pm 2.829^{e-1}$ | $1.40 \pm 1.449^{e-1}$ | 0.7276 |
| *N*-stearoyl valine | $4.868^{e-4} \pm 3.845^{e-4}$ | $5.914^{e-4} \pm 1.274^{e-4}$ | 0.5034 | $PGF_{2\alpha}$ | $1.103 \pm 1.962^{e-1}$ | $1.005 \pm 1.148^{e-1}$ | 0.6646 |
| *N*-oleoyl valine | BDL | BDL | | 6-keto$PGF_{1\alpha}$ | $3.898^{e-2} \pm 9.96^{e-3}$ | $3.614^{e-2} \pm 2.378^{e-3}$ | 0.7658 |
| *N*-nervonoyl valine | BDL | BDL | | | | | |
| *N*-linoleoyl valine | BDL | BDL | | | | | |
| *N*-docosahexaenoyl valine | BDL | BDL | | | | | |
